# ER-directed TREX1 limits cGAS recognition of micronuclei

**DOI:** 10.1101/2020.05.18.102103

**Authors:** Lisa Mohr, Eléonore Toufektchan, Kevan Chu, John Maciejowski

## Abstract

Chromosomal instability in cancer results in the formation of nuclear aberrations termed micronuclei. Spontaneous loss of micronuclear envelope integrity exposes DNA to the cytoplasm, leading to chromosome fragmentation and innate immune activation. Despite connections to cancer genome evolution and anti-tumor immunity, the mechanisms underlying damage and immune sensing of micronuclear DNA are poorly understood. Here, we use a novel method for the purification of micronuclei and live-cell imaging to show that the ER-associated nuclease TREX1 inhibits cGAS sensing of micronuclei by stably associating with and degrading micronuclear DNA upon micronuclear envelope rupture. We identify a *TREX1* mutation, previously associated with autoimmune disease, that untethers TREX1 from the ER, disrupts TREX1 localization to micronuclei, alleviates micronuclear DNA damage, and enhances cGAS recognition of micronuclei. Together, these results establish ER-directed resection of micronuclear DNA by TREX1 as a critical regulator of cytosolic DNA sensing in chromosomally unstable cells and provide a mechanistic basis for the importance of TREX1 ER-tethering in preventing autoimmunity.

## INTRODUCTION

Chromosomal instability is a long-recognized feature of cancer that is characterized by elevated rates of chromosome mis-segregation during mitosis (Bakhoum and Cantley, 2018; Maciejowski and Hatch, 2020). Chromosome mis-segregation can lead to an abnormal number of chromosomes in daughter cells or generate nuclear aberrations such as micronuclei (MN) and DNA bridges (Crasta et al., 2012; Maciejowski et al., 2015; Maciejowski and Hatch, 2020).

MN form nuclear envelopes (NEs) during telophase, but defects in assembly often result in spontaneous micronuclear envelope rupture and loss of compartmentalization during interphase (Hatch et al., 2013; Liu et al., 2018). MN with ruptured NEs (i.e. ruptured MN) exhibit compromised nuclear functions, including defective transcription and DNA replication (Crasta et al., 2012; Hatch et al., 2013; Willan et al., 2019; Zhang et al., 2015). NE rupture at MN is also associated with invasion of endoplasmic reticulum (ER) tubules into micronuclear chromatin, induction of DNA double-strand breaks (DSBs), and generation of single-stranded DNA (ssDNA) (Crasta et al., 2012; Hatch et al., 2013; Liu et al., 2018; Vietri et al. 2019; Willan et al., 2019; Zhang et al., 2015). Micronuclear DNA damage is a proposed intermediate in chromothripsis, or chromosome shattering, but the initial cause(s) of this damage are unknown (Crasta et al., 2012; Ly et al., 2017, 2019; Umbreit et al., 2020; Zhang et al., 2015).

DNA bridges are distinct nuclear aberrations that form as a result of telomere fusion or merotelic kinetochore attachment (Liu et al., 2018; Maciejowski et al., 2015; Maciejowski et al., 2019; Mackenzie et al., 2017; Umbreit et al., 2020). NE ruptures also occur at DNA bridges, but the causes are unknown (Maciejowski et al., 2015; Mackenzie et al., 2017).

NE rupture at MN enables the cytosolic DNA sensor cyclic GMP-AMP synthase (cGAS) to access chromosomal DNA (Harding et al., 2017; Mackenzie et al., 2017). Engagement with double-stranded DNA (dsDNA) stimulates cGAS catalytic activity and the production of 2′3′-cyclic GMP–AMP (cGAMP) (Ablasser et al., 2013; Civril et al., 2013; Diner et al., 2013; Gao et al., 2013; Sun et al., 2013). 2′3′-cGAMP engages with Stimulator of Interferon Genes (STING), resulting in activation of TANK Binding Kinase 1 (TBK1) and the phosphorylation and nuclear translocation of the transcription factors IRF3 and NF-κB (Ishikawa and Barber, 2008). IRF3 and NF-κB induce the expression of type I interferons and other immunomodulatory proteins (Ablasser and Chen, 2019). cGAS-STING pathway activation results in context-dependent responses that can promote or dampen anti-tumor immune responses to chromosomally unstable cancer cells (Bakhoum et al., 2018; Kwon and Bakhoum, 2020; Wang et al., 2017).

Current models of cGAS activation in ruptured MN are consistent with its biochemical preferences. cGAS recognizes dsDNA through sequence-independent interactions (Civril et al., 2013; Gao et al., 2013; Sun et al., 2013). Chromatin is capable of activating cGAS, albeit at a reduced apparent catalysis rate compared to nucleosome-free DNA (Mackenzie et al., 2017; Zierhut et al., 2019). Human cGAS is dependent on long (>45 bp) dsDNA for robust activation; ssDNA and small dsDNA fragments do not elicit robust cGAS activity (Civril et al., 2013; Kranzusch et al., 2013; Zhou et al., 2018).

TREX1 is a ubiquitously expressed, ER-associated, DNA 3′-5′ exonuclease that degrades cytosolic DNA to prevent chronic cGAS activation and consequent autoimmunity (Ablasser et al., 2014; Gray et al., 2015; Mazur and Perrino, 2001; Stetson et al., 2008; Wolf et al., 2016). Frameshift mutations that eliminate TREX1’s highly conserved C-terminal extension are associated with systemic lupus erythematosus (Lee-Kirsch et al., 2007; Richards et al., 2007). These mutations compromise TREX1 ER association, but do not affect its catalytic activity (Lee-Kirsch et al., 2007). Previous characterization of TREX1 substrates have included cytosolic DNAs derived from endogenous retroelements and byproducts of DNA metabolism (De Cecco et al., 2019; Stetson et al., 2008). We and others have previously shown that TREX1 localizes to and resects the chromosomal DNA isolated in DNA bridges, but the consequences of this for innate immune activation, including any potential impact on cytosolic DNA sensing, were unclear (Fouquerel et al., 2019; Maciejowski et al., 2015; Maciejowski et al., 2019; Xia et al., 2019). Likewise, the functional significance of TREX1’s association with the ER is also unknown.

Here, we report the discovery that TREX1 limits cGAS recognition of MN by partially degrading chromosomal DNA in ruptured MN. Using a novel method for purifying MN, we discovered that TREX1 stably associates with ruptured MN where it damages exposed micronuclear DNA. Live-cell imaging revealed that TREX1 rapidly accumulates in MN after micronuclear envelope rupture and targets micronuclear DNA for resection. TREX1 deficiency in chromosomally unstable cells causes enhanced cGAS recognition of MN, resulting in increased 2′3′-cGAMP production, IRF3 phosphorylation, and Type I interferon (IFN) induction. We further show TREX1 tethering to the ER is critical for activity in chromosomally unstable cells: a frameshift mutation in *TREX1*, previously associated with systemic lupus erythematosus (Lee-Kirsch et al., 2007; Richards et al., 2007), truncates the C-terminal region (CTR) of the protein, disrupts TREX1 association with the ER, compromises TREX1 activity in MN, and relieves cGAS inhibition. Analysis of a chimeric protein wherein the TREX1 CTR was replaced with the transmembrane domain of the core ER transporter protein Sec61 revealed that ER-tethering is critical for TREX1 activity and cGAS inhibition at ruptured MN. Together, our results identify TREX1 as a critical regulator of immune sensing in chromosomally unstable cells, define new disease-relevant substrates of TREX1, and establish ER-tethering as a critical director of TREX1 nucleolytic activity.

## RESULTS

### Purification of ruptured MN reveals association with TREX1 and evidence of ssDNA

To determine why micronuclear envelope rupture is associated with DNA damage, we developed a strategy to purify MN with ruptured NEs. MN can be separated from primary nuclei by differential centrifugation, but this method does not differentiate between ruptured and intact MN (Ly et al., 2017; Shimizu et al., 1996). Our strategy uses genetically-encoded markers and flow cytometry to separate ruptured MN from intact MN and primary nuclei (Figure 1A). To this end, HEK293T cells were labeled with H2B-mCherry and GFP-cGAS to mark chromatin and ruptured MN, respectively (Figure 1B). GFP-cGAS is a reliable marker for ruptured MN as it localizes to ruptured MN within minutes of NE disruption (Harding et al., 2017; Mackenzie et al., 2017).

**Figure 1.**
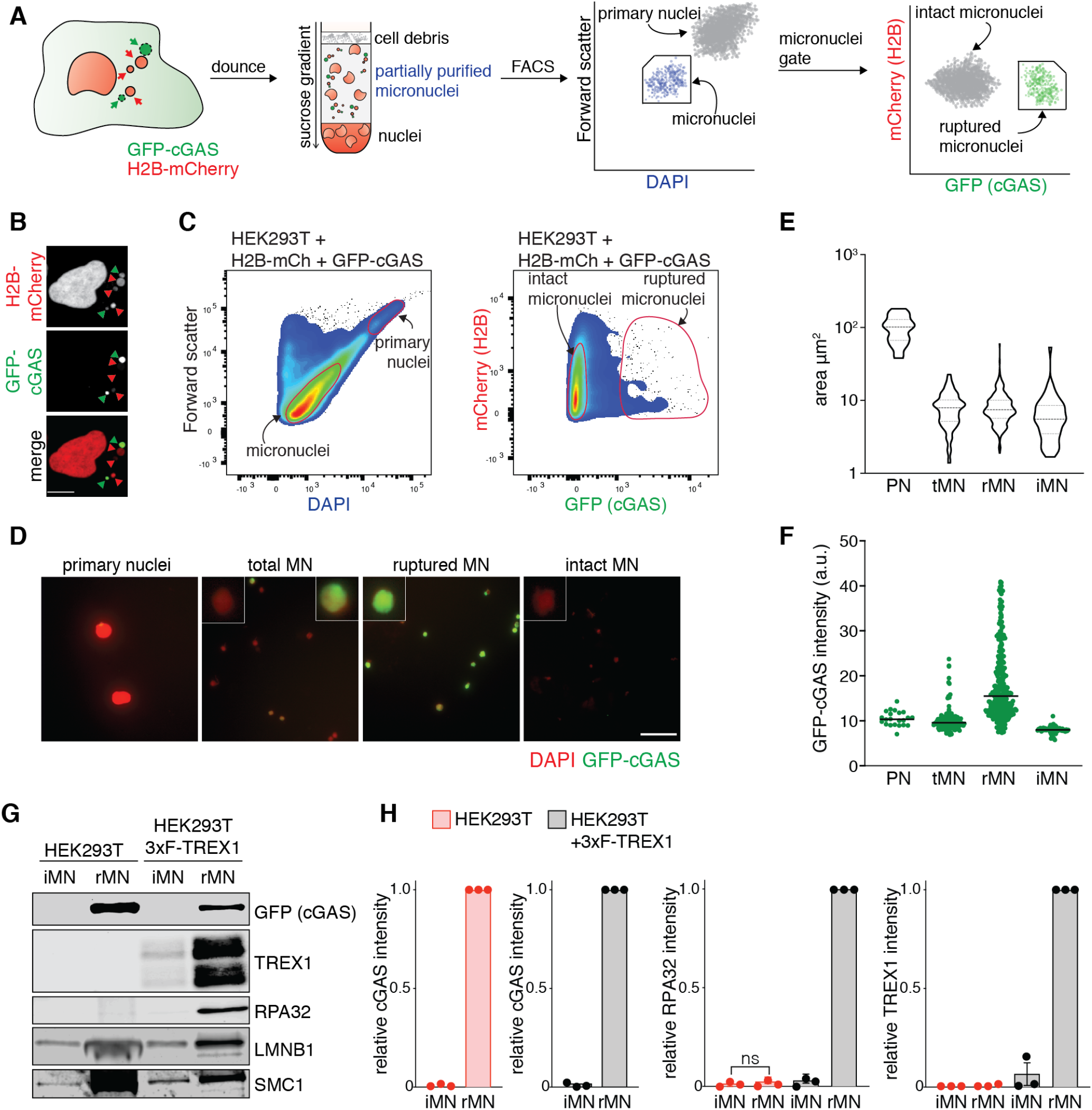
Purification of intact and ruptured MN. (A) Schematic of the purification of intact and ruptured MN. HEK293T cells expressing GFP-cGAS and H2B-mCherry as markers of ruptured MN and chromatin, respectively, were treated with Mps1i for 48 h to induce MN formation. Cells were lysed by Dounce homogenization, and lysates were processed through a sucrose density gradient centrifugation to separate MN from primary nuclei. DAPI staining, as well as GFP and mCherry fluorescent signals, were used to sort ruptured from intact MN by FACS. (B) Immunofluorescence for mCherry and GFP (cGAS) in Mps1i-treated HEK293T (+H2B-mCherry +GFP-cGAS) cells. GFP (cGAS) staining serves as a marker for ruptured MN. Green arrowheads mark ruptured MN, red arrowheads mark intact MN. Scale bar = 10 μm. (C) Flow profiles of intact and ruptured MN isolated by flow sorting. (D) Immunofluorescence staining for GFP (cGAS) and DAPI staining of DNA of the indicated fractions after FACS. Scale bar = 25 μm. (E) Measurement of the area of sorted primary nuclei (PN) and the different fractions sorted by flow cytometry. Violin plot of the areas of 22 PN, 139 tMN, 342 rMN and 54 iMN. (F) Quantification of the GFP-cGAS signal intensity as shown in (D). Signal intensity with median of 22 PN, 139 tMN, 342 rMN and 54 iMN are shown. (G) Immunoblotting for the indicated proteins in ruptured and intact MN sorted from parental and 3xFLAG-TREX1 overexpressing HEK293T (+H2B-mCherry +GFP-cGAS) cells. (H) Quantification of relative cGAS, TREX1 and RPA32 signals normalized to loading control (SMC1 or histone H3) as shown in (G). The cGAS signal was independently normalized in each cell line to correct for differences in GFP-cGAS transgene expression. RPA32 and TREX1 signals were transformed as fractions of the peak value in each experiment. Mean and s.d. of *n* = 3 independent biological replicates are shown. PN: primary nuclei, tMN: total MN, iMN: intact MN, rMN: ruptured MN.

To purify ruptured MN, HEK293T cells were treated with the Mps1 inhibitor reversine 48 h before lysis to induce chromosome mis-segregation and generate MN (Santaguida et al., 2010). Homogenates were separated by density gradient centrifugation to partially remove cell debris and primary nuclei (Figure 1A). Flow cytometric analysis of the MN-containing fractions revealed two populations that could be distinguished by distinct forward scatter and DAPI-staining intensity (Figure 1C). Image analysis of flow-purified particles identified these populations as primary nuclei and MN (Figure 1D). Further inspection of the MN population by flow cytometry revealed two populations marked by differences in GFP-cGAS signal intensity (Figure 1C). Image analysis showed that all purified MN exhibited a similar distribution of surface area averaging approximately 8.5 μm2 across populations; however, the average GFP-cGAS signal intensity differed more than two-fold across the two populations, indicating that they correspond to intact and ruptured MN (Figure 1D-F).

Analysis of purified MN by immunoblotting confirmed that GFP-cGAS exhibited marked changes across the two populations (Figure 1G, Figure S1A). Additionally, the ESCRT-III subunit CHMP2A, which localizes to MN after micronuclear envelope rupture (Vietri et al., 2019; Willan et al., 2019), was also present in GFP-cGAS positive MN, indicating that this population likely corresponds to ruptured MN (Figure S1A). Taken together, these data validate the utility of this method for purifying ruptured MN.

We next determined whether purified MN showed evidence of DNA damage after micronuclear envelope rupture. Surprisingly, neither RPA32 nor γH2AX were present at increased levels in ruptured MN (Figure 1G, lanes 1 and 2, Figure S1B and S1C). Based on our previous work (Maciejowski et al., 2015), we reasoned that the absence of micronuclear DNA damage could be explained by lack of TREX1, which is not expressed in HEK293T cells (Figure S1D). To test this, HEK293T cells were modified to express 3×FLAG-TREX1 (Figure S1D). Immunoblotting analysis of ruptured MN purified from TREX1-expressing HEK293T cells showed an approximately 50-fold increase in RPA32 relative to intact MN indicating the presence of TREX1-dependent ssDNA in ruptured MN (Figure 1G and 1H, Figure S1B). TREX1 expression did not affect γH2AX levels in ruptured MN suggesting additional sources of micronuclear DNA damage (Figure S1B and S1C). Surprisingly, given its association with the ER, we also found that TREX1 co-purified with ruptured MN, indicating stable association of TREX1 with purified, ruptured MN (Figure 1G and 1H). These data suggest that TREX1 may damage and stably associate with chromatin after micronuclear envelope rupture.

### TREX1 rapidly accumulates in ruptured MN

To better understand the observed association of TREX1 with purified, ruptured MN, we analyzed TREX1 localization in chromosomally unstable cells. We previously observed that TREX1 localized to MN and DNA bridges, but the dynamics of TREX1 targeting were not investigated (Maciejowski et al., 2015). Live-cell imaging showed that GFP-TREX1, expressed in MCF10A *TREX1* KO cells, displayed perinuclear localization consistent with its ER association (Figure 2A; Figure S2A) (Stetson et al., 2008). To investigate TREX1 localization to MN, we treated non-tumorigenic, chromosomally stable MCF10A cells with reversine. Live-cell imaging showed that GFP-TREX1 localized to approximately 40% of MN within 24 hours of mitotic exit (Figure 2A and 2B). Analysis of GFP-TREX1 in a panel of chromosomally unstable breast cancer cell lines revealed similar patterns of GFP-TREX1 localization to MN (Figure S2B).

**Figure 2.**
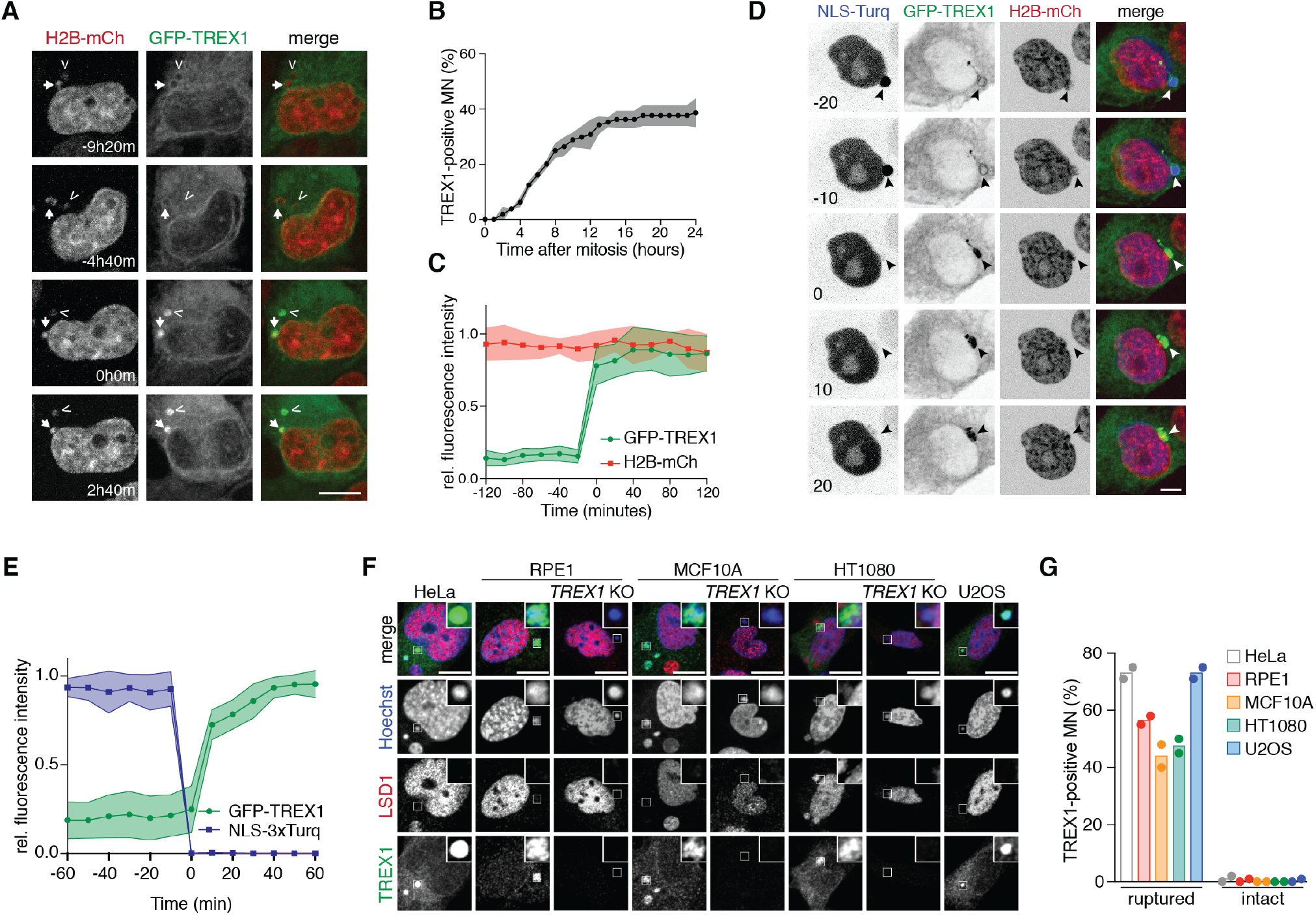
TREX1 localizes to MN after micronuclear envelope rupture. (A) Frames from live-cell imaging of Mps1i-treated MCF10A cells expressing GFP-TREX1 and H2B-mCherry to mark chromatin. Time relative to TREX1 localization to MN is marked. Arrowhead and arrow mark MN. (B) Quantification of the cumulative percentage of TREX1-positive MN as shown in (A). MN were followed from mitotic exit until the subsequent mitosis or for a maximum of 24 h. Error bars show s.d. of *n* = 45 MN quantified across 3 independent biological replicates. (C) Quantification of the relative fluorescence signal intensities of GFP-TREX1 and H2B-mCherry at MN (t = 0, initial TREX1 localization) as shown in (A). Relative mean fluorescence intensities and s.d. of *n* = 11 MN are shown. (D) Frames from live-cell imaging of reversine-treated MCF10A cells expressing GFP-TREX1, H2B-mCherry and NLS-3×mTurquoise2 to assay for NE rupture (t = 0, micronuclear envelope rupture). Quantification of the relative fluorescence signal intensities of GFP-TREX1 and NLS-3×mTurquoise2 as shown in (D). Mean and s.d. of *n* = 15 MN are shown. Immunofluorescence for TREX1 and LSD1 in the indicated cell lines. LSD1 staining serves as a marker for intact MN (Hatch et al., 2013). DNA was stained with Hoechst 33342. Insets show magnified regions (marked by box) to highlight TREX1 localization to the MN. (G) Quantification of the percentage of TREX1-positive ruptured and intact MN as shown in (F). Mean of *n* =2 independent biological replicates are shown (>90 MN quantified per replicate and cell line). All scale bars = 10 μm.

Based on prior studies and our data showing the presence of TREX1 in ruptured, but not intact, purified M N, w e r easoned t hat T REX1 m ay localize to MN after micronuclear envelope rupture (Harding et al., 2017; Hatch et al., 2013; Mackenzie et al., 2017) (Figure 1G and 1H). To test this, we assayed for micronuclear envelope rupture using a previously described NLS-3×mTurquoise2 marker (Hatch et al., 2013; Maciejowski et al., 2015; Vargas et al., 2012). This analysis showed that GFP-TREX1 localization to MN was coincident with micronuclear envelope rupture, as measured by loss of NLS-3×mTurquoise2 from MN (Figure 2D and 2E). In addition, analysis of micronuclear envelope rupture using mTurquoise2-tagged cGAS showed that TREX1 and cGAS localize to MN with comparable timing (Figure S2C and S2D). Immunofluorescence analysis of a panel of cell lines confirmed t he p resence of endogenous TREX1 at 40%-70% of ruptured MN and less than 5% of intact MN (Figure 2F and 2G). Likewise, transgenic TREX1 was present at 70% of ruptured MN in HEK293T cells (Figure S2E and S2F). *TREX1* deletion in RPE1, MCF10A, and HT1080 cells validated the specificity o f o ur a ntibody (Figure 2F and 2G; Figure S2A). Based on these data, we conclude that TREX1 localizes to MN following micronuclear envelope rupture.

### ER-tethering is necessary for targeting TREX1 to ruptured MN

We next sought to understand how TREX1 is targeted to MN after micronuclear envelope rupture. TREX1 has high affinity for ssDNA and dsDNA and possesses a highly conserved, single-pass transmembrane helix at its C-terminus that anchors TREX1 in the ER and positions the nuclease domain in the cytosol (Lee-Kirsch et al., 2007; Mazur and Perrino, 2001; Wolf et al., 2016). Deleting this C-terminal extension compromises TREX1 association with the ER but does not affect its catalytic activity (Lee-Kirsch et al., 2007; de Silva et al., 2007). Nevertheless, this C-terminal extension appears to be critical for TREX1 function, as frameshift mutations that truncate the protein are associated with systemic lupus erythematosus (Lee-Kirsch et al., 2007; Richards et al., 2007).

Micronuclear envelope rupture is associated with the invasion of ER tubules into the disrupted chromatin of MN (Hatch et al., 2013; Willan et al., 2019). Interestingly, we observed that the ER lumen protein Calreticulin was always present in TREX1-positive MN (Figure 3A and 3B), suggesting that TREX1 localization to ruptured MN may depend on its ER association. To test this, we deleted the C-terminal 79 residues that are unique to TREX1 (TREX1ΔC) and not found in the otherwise highly homologous TREX2 (Figure 3C). As previously reported (Lee-Kirsch et al., 2007; Stetson et al., 2008), GFP-TREX1ΔC was diffusively localized throughout the cell, indicating apparent dissociation from the ER (Figure 3D and 3E, Figure S3B). GFP-TREX1 was present at approximately 75% of ruptured MN, while GFP-TREX1ΔC was never present at MN, even after micronuclear envelope rupture, indicating a severe defect in normal localization (Figure 3E and 3F, Figure S3B and S3C).

**Figure 3.**
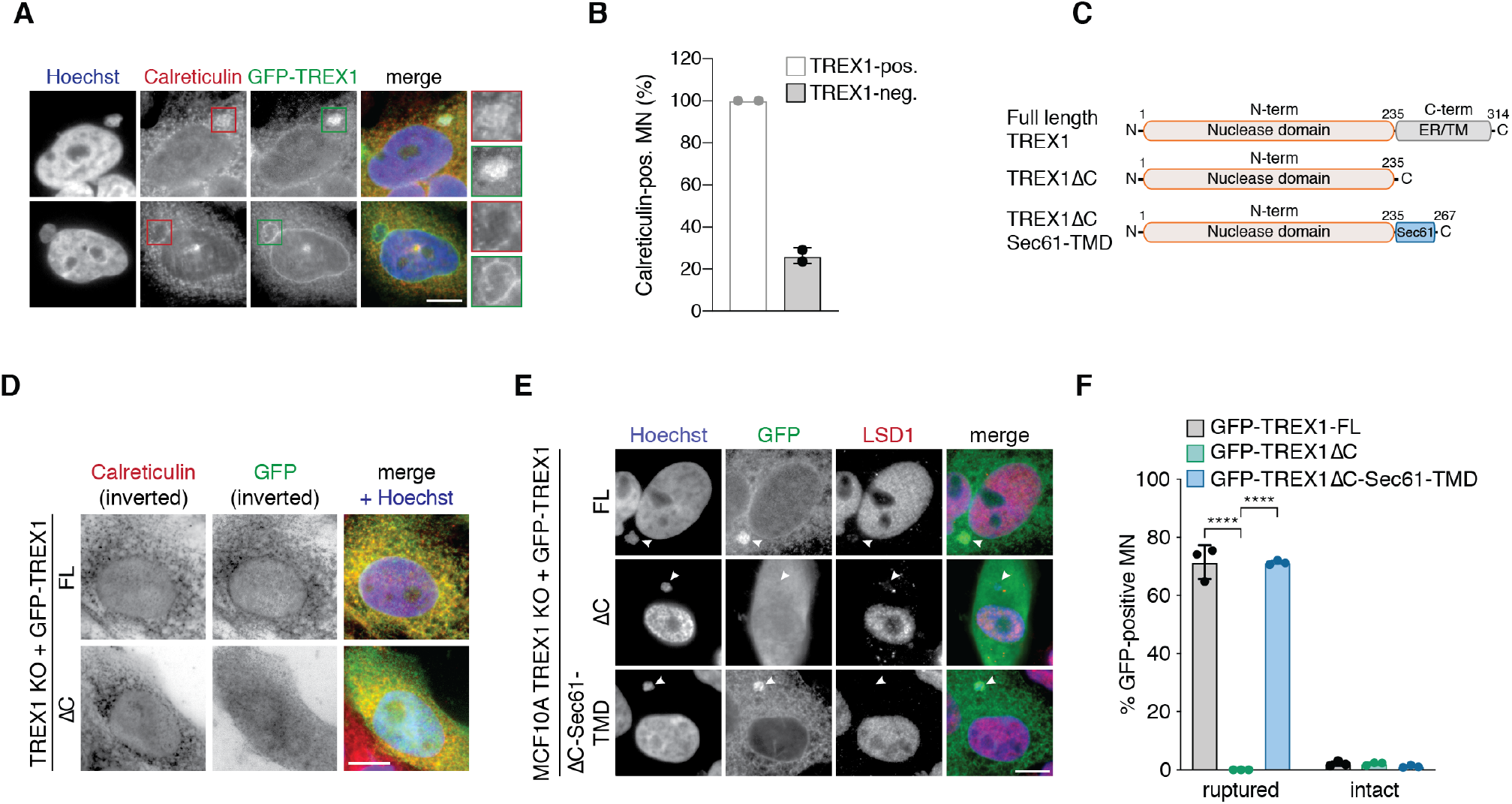
ER-tethering of TREX1 mediates recruitment to ruptured MN. (A) Immunofluorescence for GFP (TREX1) and Calreticulin (ER marker) in Mps1i-treated MCF10A *TREX1* KO 1 cells expressing GFP-TREX1. Top row shows ER-invaded MN (Calreticulin-positive), bottom row shows Calreticulin-negative MN. Magnified MN are shown on the right (red frame: Calreticulin, green frame: GFP). (B) Quantification of the percentage of Calreticulin-positive MN within TREX1-positive and -negative MN as shown in (A). Mean and s.d. of *n* = 2 independent biological replicates are shown (>135 MN quantified per replicate and cell line). (C) Schematic of full length TREX1, TREX1ΔC, and TREX1ΔC-Sec61-TMD, a chimeric protein consisting of TREX1ΔC fused to the Sec61 transmembrane domain. (D,E) Immunofluorescence for GFP (TREX1) and (D) Calreticulin (ER marker) or (E) LSD1 in the indicated cell lines. DNA was stained with Hoechst 33342. Arrowheads mark ruptured MN. (F) Quantification of the percentage of GFP-TREX1-positive ruptured and intact MN as shown in (E). Mean and s.d. of *n* = 3 independent biological replicates are shown (>180 MN quantified per replicate and cell line). *P* values were calculated by one-way ANOVA with Tukey’s multiple comparisons test (\*\*\*\**P* < 0.0001). All scale bars = 10 μm.

To determine if loss of ER-tethering could explain the TREX1ΔC localization defect, we re-tethered TREX1ΔC to the ER via fusion to the transmembrane domain of Sec61 (hereafter TREX1ΔC-Sec61-TMD), a component of the protein-conducting channel that mediates protein transport across ER membranes (Figure 3C, Figure S3A). Similar to GFP-TREX1, GFP-TREX1ΔC-Sec61-TMD localized to 75% of ruptured MN (Figure 3E and 3F). Taken together, these data indicate that TREX1 localization to ruptured MN is dependent on its association with the ER.

### TREX1 resects DNA in ruptured MN

We next sought to understand the functional significance of TREX1 localization to ruptured MN. The presence of RPA32 in ruptured MN purified from TREX1-proficient HEK293T cells suggested that TREX1 may resect DNA in ruptured MN (Figure 1F and 1G). Indeed, live-cell imaging showed immediate GFP-RPA70 accumulation in MN after micronuclear envelope rupture indicating the presence of ssDNA (Figure 4A). Additionally, we quantified t he presence of RPA32 foci in ruptured and intact MN using LSD1 staining as a marker for intact MN as previously described (Hatch et al., 2013). RPA32 foci were visible in 40% of ruptured MN compared with less than 5% of intact MN (Figure 4B and 4C). Similar to our prior observations at DNA bridges (Maciejowski et al., 2015), two independently derived *TREX1* KO cell lines showed nearly complete loss of RPA32 accumulation in ruptured MN (Figure 4B and 4C). In agreement with our immunoblotting analyses of purified M N, RPA32 foci were present in nearly 60% of ruptured MN in TREX1-proficient H EK293T c ells b ut l ess t han 10% of ruptured MN in TREX1-deficient H EK293T cells (Figure S4A and S4B).

**Figure 4.**
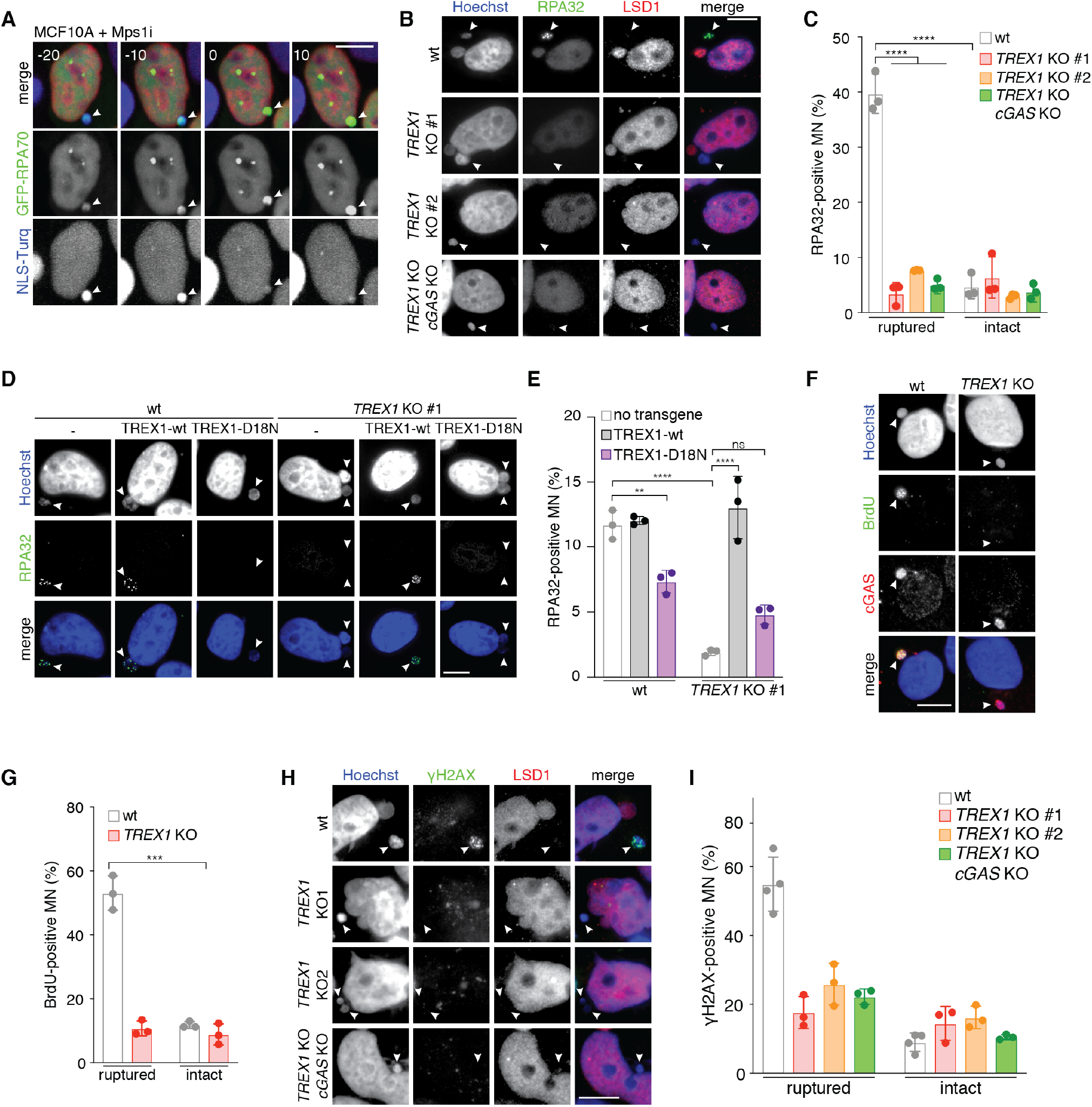
TREX1 degrades micronuclear DNA after micronuclear envelope rupture. (A) ssDNA accumulates in MN after micronuclear envelope rupture. Frames from live-cell imaging of Mps1i-treated MCF10A cells expressing H2B-mCherry (red), GFP-RPA70 and NLS-3×mTurquoise2 (NLS-Turq). Arrowheads mark MN. Time (minutes) is relative to micronuclear envelope rupture (t = 0, micronuclear envelope rupture). (B-E) Accumulation of ssDNA in MN is dependent on TREX1. Indicated MCF10A cells were treated with Mps1i for 72 h. (B) Immunofluorescence for RPA32 and LSD1, DNA was stained with Hoechst 33342. Arrowheads mark ruptured MN. Loss of LSD1 signal was used to infer micronuclear envelope rupture. (C) Quantification of the percentage of ruptured and intact MN positive for RPA32 foci as shown in (B). Mean and s.d. from *n* = 3 independent biological replicates are shown (>200 MN quantified per replicate and cell line). *P* values were calculated by two-way ANOVA with Sidak’s multiple comparisons test (\*\*\*\**P* < 0.0001). (D) Immunofluorescence for RPA32, DNA was stained with Hoechst 33342. Arrowheads mark MN. (E) Quantification of the percentage of RPA32-positive MN as shown in (D). Mean and s.d. from *n* = 3 independent biological replicates are shown (>220 MN quantified per replicate and cell line). *P* values were calculated by two-way ANOVA with Sidak’s multiple comparisons test (\*\*\*\**P* < 0.0001). (F) Detection of ssDNA in MN after Mps1i treatment. Cells were labeled with BrdU for 24 h, before treatment with Mps1i for 48 h. Immunofluorescence for BrdU (ssDNA) and cGAS (ruptured MN), DNA was stained with Hoechst 33342. Arrowheads mark ruptured MN. (G) Quantification of the percentage of BrdU-positive ruptured and intact MN as shown in (F). Mean and s.d. from *n* = 3 independent biological replicates are shown (>180 MN quantified per replicate and cell line). *P* value was calculated by Student’s t-test (\*\*\**P* = 0.0002). (H) Immunofluorescence for γH2AX and LSD1 in the indicated MCF10A cells. DNA was stained with Hoechst 33342. Arrowheads mark ruptured MN. (I) Quantification of the percentage of ruptured and intact MN with detectable γH2AX foci as shown in (H). Mean and s.d. from *n* = 3 independent biological replicates are shown (>180 MN quantified per replicate and cell line). *P* values were calculated by two-way ANOVA with Sidak’s multiple comparisons test (\*\*\*\**P* < 0.001, ns = not significant). All scale bars = 10 μm.

Reintroduction of wild-type TREX1, but not a catalytically-deficient m utant (TREX1-D18N) (Lehtinen et al., 2008), restored RPA32 accumulation in MN (Figure 4D and 4E, Figure S4C). Consistent with prior reports (Lehtinen et al., 2008; Maciejowski et al., 2015; Xia et al., 2019), TREX1-D18N had a dominant-negative effect in the TREX1-proficient cell line, significantly reducing the appearance of RPA32 in MN (Figure 4D and 4E).

In a complementary approach, we monitored ssDNA formation in MN by quantifying 5-bromo-2’-deoxyuridine (BrdU) levels in MN under non-denaturing conditions (Figure 4F and 4G). This assay is based on the principle that incorporated BrdU cannot be detected under native conditions unless exposed by resection (Tkáč et al., 2016). This analysis showed that the frequency of BrdU-positive ruptured MN was reduced 5-fold in *TREX1* KO cells, further supporting a model of TREX1-dependent DNA resection in ruptured MN (Figure 4F and 4G).

γH2AX staining in ruptured MN was also reduced in *TREX1* KO and TREX1-D18N reconstituted cells, albeit to a lesser extent than ssDNA (Figure 4H and 4I, Figure S4D and S4E). In contrast, TREX1 expression in HEK293T cells did not result in a measurable increase in γH2AX signal in ruptured MN, indicating additional sources of DNA damage in HEK293T cells (Figure S4F and S4G). Importantly, *TREX1 cGAS* double KO cells exhibited similar defects in the levels of ssDNA and γH2AX, suggesting that the observed defects reflected l ack o f TREX1 catalytic a ctivity r ather than potential, indirect consequences of increased cGAS activation (Figure 4B and 4C, Figure 4H and 4I). Collectively, these data suggest that TREX1 resects DNA in ruptured MN to generate ssDNA.

### TREX1 limits cGAS recognition of ruptured MN and DNA bridges

We reasoned that TREX1 resection of micronuclear DNA may suppress cGAS surveillance of ruptured MN. To test this, we treated MCF10A cells with reversine to induce chromosome mis-segregation and generate MN and DNA bridges (Figure 5A and 5B, Figure S5A and S5B) (Maciejowski et al., 2010, 2015). cGAS signal intensity and localization to MN was similar in wild-type and *TREX1* KO cells (Figure 5C and 5D, Figure S5C and S5D). To assess cGAS activation at MN we measured 2′3′-cGAMP production in Mps1-inhibited cells by ELISA (Figure 5E). This analysis showed that wild-type MCF10A cells accumulated less than 20 fmol of 2′3′-cGAMP per million cells over 3 days of Mps1 inhibition (Figure 5E). In contrast, *TREX1* KO cells accumulated approximately 80–120 fmol of 2′3′-cGAMP per million cells, a significant increase over wild-type MCF10A cells (Figure 5E). 2′3′-cGAMP production was also increased after Mps1 inhibition in TREX1 hypomorphic cells (Figure 5E). *cGAS* deletion completely blocked 2′3′-cGAMP production, confirming the specificity of our assay (Figure 5E).

**Figure 5.**
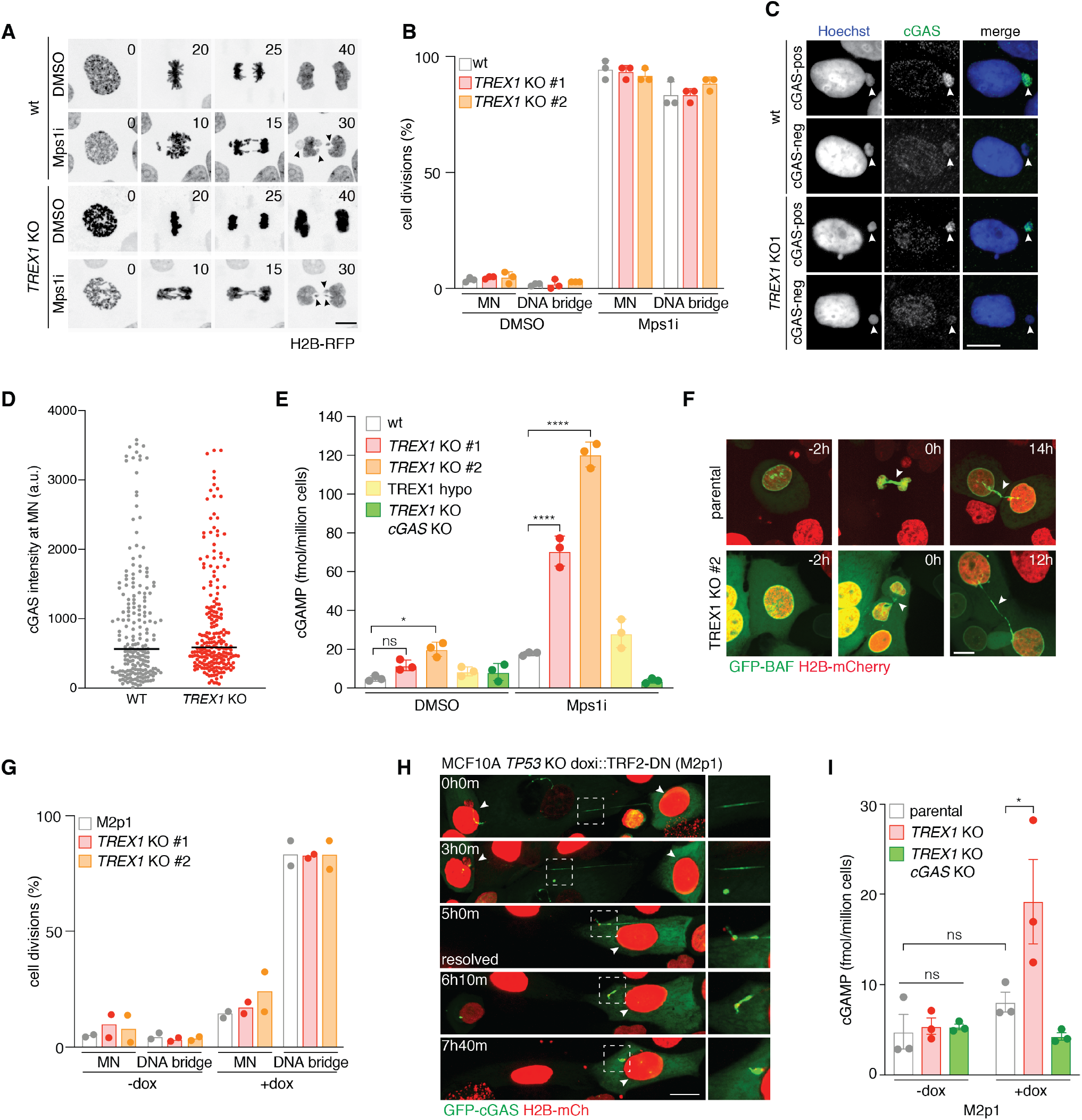
TREX1 inhibits cGAS activation at MN and DNA bridges. (A) Extracted frames from live-cell imaging of DMSO- or reversine-treated MCF10A cells of the indicated genotypes expressing H2B-RFP to mark chromatin. Insets mark relative time in minutes since mitotic NE breakdown (t = 0 minutes). Arrowheads mark mis-segregating chromosomes. (B) Quantification of cell divisions resulting in the formation of MN or DNA bridges as a percentage of divisions as shown in (A). Mean and s.d. of *n* = 3 independent biological replicates (>50 cells analyzed per experiment) are shown. (C) Immunofluorescence staining for cGAS and Hoechst staining of DNA of the indicated MCF10A cells 72 h after treatment with Mps1i. (D) Quantification of cGAS signal intensity as shown in (C). Signal intensity with median of *n* = 3 independent biological replicates are shown (>60 MN quantified per experiment). (E) TREX1 inhibits cGAS activation at MN. ELISA analysis of 2 3-cGAMP production in the indicated cells 72 h after treatment with Mps1i. Mean and s.d. of *n* = 3 independent biological replicates are shown. *P* values were calculated by one-way ANOVA with Tukey’s multiple comparisons test (\**P* < 0.05, \*\**P* < 0.01, \*\*\*\**P* < 0.0001, ns = not significant). (F) TREX1 deficiency does not alter DNA bridge formation in M2p1 cells. Frames from live-cell imaging of doxycycline-treated M2p1 (MCF10A *TP53* KO doxi::TRF2-DN) cells expressing GFP-BAF to mark DNA bridges and H2B-mCherry to mark chromatin. Relative time between images is marked in hours in the top right of each panel. Arrowheads mark DNA bridges. Scale bar = 10 μm. (G) Quantification of the number of cell divisions resulting in MN or DNA bridge formation (F). Mean of *n* = 2 independent biological replicates (>50 cells analyzed per condition) is shown. (H) Frames from live-cell imaging of doxycycline-treated M2p1 (MCF10A *TP53* KO doxi::TRF2-DN) cells expressing GFP-cGAS and H2B-mCherry to mark chromatin. Elapsed time is marked in hours and minutes in the top left of each panel. Arrowheads mark nuclei connected by DNA bridge. Insets show magnified regions (marked by dotted box) to highlight cGAS localization to the DNA bridge. Frame showing bridge resolution is marked. (I) TREX1 inhibits cGAS activation at DNA bridges. ELISA analysis of 2 3-cGAMP production in the indicated cells 72 h after treatment with doxycycline to induce telomere fusions and DNA bridges. Mean and s.d. of *n* = 3 independent biological replicates are shown. *P* values were calculated by one-way ANOVA with Tukey’s multiple comparisons test (\**P* < 0.05, \*\**P* < 0.01, \*\*\*\**P* < 0.0001, ns = not significant). All scale bars = 10 μm.

The increased 2′3′-cGAMP accumulation in *TREX1* KO cells after Mps1 inhibition could not be explained by altered response to Mps1 inhibition, as wild-type and *TREX1* KO cells exhibited similar numbers of mitotic errors and MN (Figure 5A and 5B, Figure S5A and S5B). Likewise, we could not detect increased MN-independent, cytosolic dsDNA levels after *TREX1* deletion or Mps1 inhibition (Figure S5E-H). cGAS exists in nuclear and cytosolic fractions, associating with chromatin following mitotic NE breakdown and accumulating in the cytosol during the subsequent cell cycle (Gentili et al., 2019; Volkman et al., 2019; Yang et al., 2017). We reasoned that unexpected alterations in cGAS subcellular distribution may increase 2′3′-cGAMP in the *TREX1* KO cells. However, cGAS exhibited similar cytosolic and nuclear localization patterns in wild-type and *TREX1* KO cells (Figure S5I and S5J).

To further assess TREX1-mediated regulation of cGAS in chromosomally unstable cells, we adapted a previously described cell model of telomere crisis as a distinct system to induce chromosomal instability in MCF10A cells (Maciejowski and de Lange, 2017; Maciejowski et al., 2015). Briefly, we use doxycycline-inducible expression of a dominant negative allele of the Shelterin complex subunit TRF2 to generate dicentric chromosomes (Figure S6A) (van Steensel et al., 1998). *TP53* was deleted to enable cell cycle progression in the presence of persistent telomere dysfunction (Karlseder et al., 1999). Consistent with our prior study (Maciejowski et al., 2015), dicentric chromosomes induced in this system persisted through mitosis to form long DNA bridges with high penetrance (approximately 80% of cell divisions), while MN were rarely generated (Figure 5F and 5G).

Local loss of Lamin B1 signal at DNA bridges indicated likely NE rupture in wild-type and *TREX1* KO cells (Figure S6B and S6C). Indeed, GFP-TREX1 and GFP-cGAS were both present on DNA bridges and remained associated with broken DNA bridges (Figure 5H, Figure S6D-F). Despite extensive cGAS accumulation on DNA bridges, inducing telomere crisis did not result in a detectable increase in 2′3′-cGAMP production (Figure 5I). However, *TREX1* KO cells accumulated nearly 20 fmol of 2′3′-cGAMP per million cells after inducing telomere crisis, a >2-fold increase over their TREX1-proficient counterparts (Figure 5I). Once again, *cGAS* deletion completely blocked 2′3′-cGAMP production (Figure 5I). Taken together, these data suggest that TREX1 limits cGAS recognition of ruptured MN and DNA bridges.

### TREX1 suppresses cGAS-dependent inflammatory responses to chromosomal instability

We next sought to determine if enhanced cGAS recognition of MN and DNA bridges in *TREX1* KO cells impacted cGAS-STING pathway activity. Following cGAS-STING activation, TBK1 phosphorylates the transcription factor IRF3 at two residues (S386 and S396), inducing IRF3 dimerization and transcription of type I IFN (Liu et al., 2015). Surprisingly, given the frequent MN accumulation and modest increases in 2′3′-cGAMP production, IRF3 S386 phosphorylation was not elevated in wild-type MCF10A cells after Mps1 inhibition (Figure 6A and 6B). In contrast, IRF3 S386 phosphorylation was increased 3 to 4-fold in two independently derived *TREX1* KO cell lines after Mps1 inhibition (Figure 6A and 6B). Following reversine treatment, TREX1 protein levels were unchanged, indicating that the observed effects occurred in the absence of TREX1 upregulation (Figure 6A and 6C). Immunofluorescence a nalysis o f IRF3 S396 phosphorylation showed a subtle, but detectable increase in nuclear foci after Mps1 inhibition, which was significantly i ncreased i n *T REX1* KO c ells (Figure 6D and 6E).

**Figure 6.**
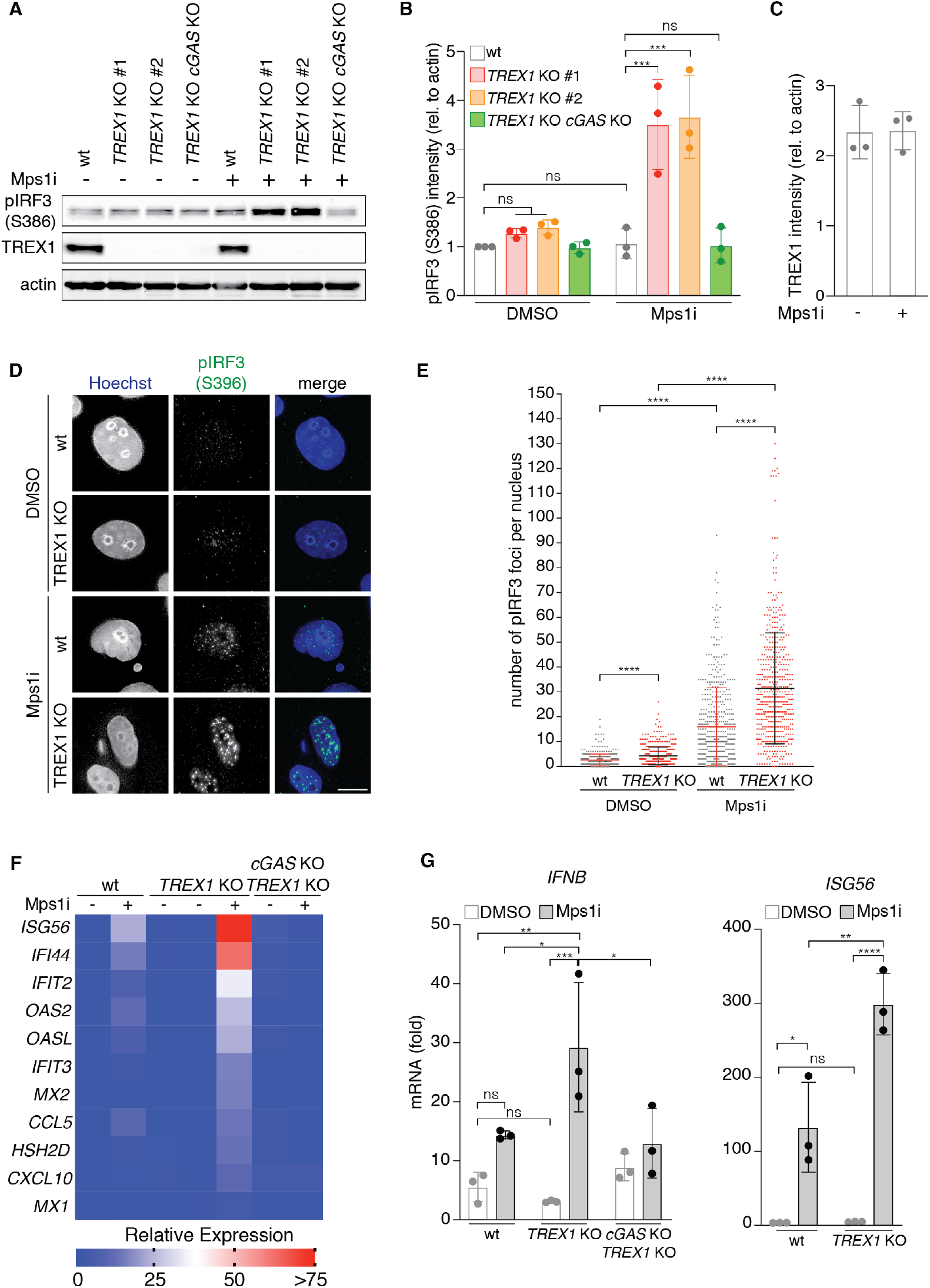
TREX1 inhibits immune sensing of chromosomal instability. (A) Immunoblotting for IRF3 S386 phosphorylation, TREX1 and actin in the indicated MCF10A cells 72 h after DMSO or Mps1i treatment. (B) Quantification of pIRF3(S386) relative to corresponding actin signal as shown in (A). Mean and s.d. of *n* = 3 independent biological replicates are shown. *P* values were calculated by one-way ANOVA with Tukey’s multiple comparisons test (\*\*\**P* < 0.001, ns = not significant). (C) Quantification of TREX1 relative to corresponding actin signal as shown in (A). Mean and s.d. of *n* = 3 independent biological replicates are shown. (D) Immunofluorescence for IRF3 S396 phosphorylation in the indicated MCF10A cells 72 h after treatment with DMSO or Mps1i. DNA was stained with Hoechst 33342. Scale bar = 10 μm. (E) Quantification of pIRF3 foci as shown in (D). Number of pIRF3 foci per nucleus, mean and s.d. of *n* = 2 independent biological replicates are shown (>250 cells quantified per experiment). *P* values were calculated by Kruskal-Wallis test for multiple comparisons (\*\*\*\**P* < 0.0001). (F) Heat map of Nanostring analysis of gene expression in the indicated MCF10A cells including two independently derived *TREX1* KOs. RNA was isolated 72 h after DMSO or Mps1i treatment and Nanostring analysis was performed using a custom CodeSet (see methods) (*n* = 1 biological replicate). Quantification of the relative expression of *IFNB* and *ISG56* in the indicated MCF10A cells 72 h after DMSO or Mps1i treatment by RT-qPCR. Mean and s.d. of *n* = 3 independent biological replicates are shown. *P* values were calculated by one-way ANOVA with Tukey’s multiple comparisons test (\**P* < 0.05, \*\**P* < 0.0, \*\*\**P* < 0.001, ns = not significant).

In line with the lack of substantial IRF3 phosphorylation, Mps1 inhibition elicited a mild transcriptional response in wild-type MCF10A cells and no significant I FN p roduction (Figure 6F and 6G). However, expression of *IFNB1, ISG56, IFI44, IFIT2,* and other Interferon-Stimulated Genes (ISGs) was elevated in *TREX1* KO cells after Mps1 inhibition (Figure 6F and 6G). Consistent with prior observations (Simpson et al., 2020), *TREX1* deletion was not sufficient to generate a major inflammatory response in untreated and therefore chromosomally stable MCF10A cells (Figure 6A and 6B, Figure 6D-6G). Deleting *cGAS* in *TREX1* KO cells abolished the observed increases in IRF3 S386 phosphorylation and gene expression, indicating cGAS as the prime sensor active in this system (Figure 6A and 6B, Figure 6F and 6G).

*TREX1* deletion did not result in detectable IRF3 S386 phosphorylation or increased ISG expression in our telomere crisis model (Figure S6G-I). The lack of IRF3 phosphorylation and ISG induction probably stems from the smaller increases in 2′3′-cGAMP production observed in *TREX1* KO cells in telomere crisis model versus Mps1 inhibition (80—120 fmol 2′3′-cGAMP per million cells vs. approximately 20 fmol 2′3′-cGAMP per million cells; Figure 5E and 5I). Weaker cGAS activation in our telomere crisis model may reflect differences in the relative amounts of cytosolic DNA in these models or tension-induced disruption of B-form DNA structure at DNA bridges (Civril et al., 2013).

Taken together, these data suggest that TREX1 inhibits innate immune responses to chromosomal instability by limiting cGAS recognition of ruptured MN and DNA bridges.

### ER-tethering directs TREX1 activity to ruptured MN

We next sought to determine if ER-tethering was necessary for TREX1-dependent regulation of cGAS in chromosomally unstable cells. To this end, we examined the activities of TREX1ΔC and TREX1ΔC-Sec61-TMD (Figure 3) at ruptured MN. As expected, given its inability to localize to ruptured MN, TREX1ΔC was unable to resect DNA in ruptured MN as measured by RPA32 accumulation (Figure 7A and 7B). TREX1ΔC was also defective at generating γH2AX in ruptured MN (Figure S7A and S7B). In contrast, introducing TREX1ΔC-Sec61-TMD into *TREX1* KO cells restored nearly normal levels of ssDNA and γH2AX signal in ruptured MN (Figure 7A and 7B; Figure S7A and S7B). These results suggest that ER-tethering is necessary to direct TREX1 catalytic activity to ruptured MN.

**Figure 7.**
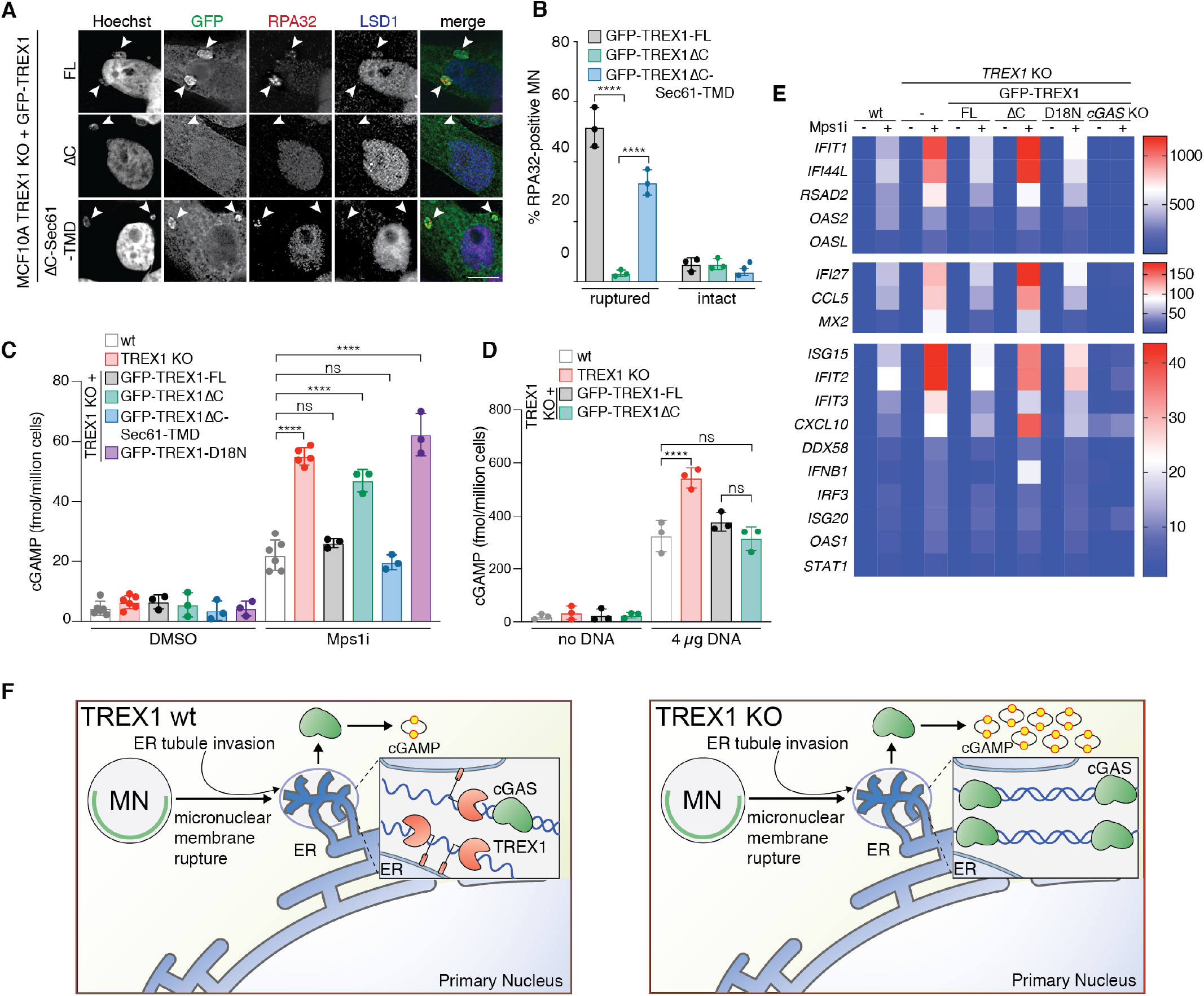
Re-tethering TREX1ΔC to the ER restores normal TREX1 function and cGAS inhibition at ruptured MN. (A) TREX1 ER-tethering is necessary for ssDNA accumulation in MN after micronuclear envelope rupture. Immunofluorescence for RPA32 and LSD1 in indicated cells treated with Mps1i for 72 h, DNA was stained with Hoechst 33342. Arrowheads mark ruptured MN. Scale bar = 10 μm. (B) Quantification of the percentage of RPA32-positive ruptured and intact MN as shown in (A). Mean and s.d. from *n* = 3 independent biological replicates are shown. *P* values were calculated by two-way ANOVA with Sidak’s multiple comparisons test (\*\*\*\**P* < 0.0001). (C,D) ELISA analysis of 2 3-cGAMP production in the indicated cells 72 h after treatment with Mps1i (C) or 24 h after dsDNA transfection (D). Mean and s.d. of *n* = 3 independent biological replicates are shown. *P* values were calculated by one-way ANOVA with Tukey’s multiple comparisons test (\*\*\*\**P* < 0.0001, ns = not significant). (E) Heat map of Nanostring analysis of gene expression in the indicated MCF10A cells 72 h after Mps1i treatment. Nanostring analysis was performed using a custom CodeSet (see methods) (*n* = 3 independent biological replicates). (F) Model of TREX1 action at ruptured MN (rMN). See discussion for details.

We next asked if TREX1 ER-tethering was required for effective cGAS inhibition. Analysis of cGAS activity via 2′3′-cGAMP ELISA showed that *TREX1* KO and TREX1 D18N and TREX1ΔC-reconstituted cells accumulated 50—60 fmol of 2′3′-cGAMP per million cells, indicating that TREX1ΔC failed to suppress cGAS activation in chromosomally unstable cells (Figure 7C). In contrast, *TREX1* KO cells reconstituted with full-length TREX1 or TREX1ΔC-Sec61-TMD accumulated 20—25 fmol of 2′3′-cGAMP after Mps1 inhibition (Figure 7C). These data indicate that TREX1 tethering to the ER is critical for cGAS inhibition in chromosomally unstable cells.

To test whether ER-tethering is required for the TREX1-dependent regulation of cGAS sensing of nucleosome-free DNA, we measured 2′3′-cGAMP production after plasmid DNA transfection. TREX1ΔC-reconstituted cells behaved similarly to cells reconstituted with wild-type TREX1 accumulating approximately 300 fmol of 2′3′-cGAMP per million cells after plasmid DNA transfection, while *TREX1* KO cells contained nearly 600 fmol of 2′3′-cGAMP per million cells (Figure 7D). Nevertheless, we do not rule out the possibility that ER-tethering may promote TREX1 engagement with transfected DNA, which has previously been shown to associate with ER membranes (Wang et al., 2016). We conclude that TREX1 tethering to the ER is critical for micronuclear DNA degradation and for suppressing cGAS activity in chromosomally unstable cells.

Gene expression analysis further highlighted the necessity of ER-tethering for effective inhibition of the cGAS-STING pathway in chromosomally unstable cells (Figure 7E). Similar to our initial results (Figure 6F and 6G), Mps1 inhibition elicited a mild transcriptional response, which included the cGAS-dependent upregulation of pro-inflammatory, anti-viral transcripts such as *IFIT1, IFI44L, RSAD2, OAS2, ISG15*, and others (Figure 7E). Transcription of these targets was further enhanced in *TREX1* KO cells. Notably, *IFNB1* expression, which was increased 2.9-fold in wild-type cells after Mps1 inhibition, was increased 8.3-fold in Mps1-inhibited *TREX1* KO cells (Figure 7E). Reintroducing full length TREX1, but not TREX1ΔC or the nuclease deficient TREX1-D18N, into *TREX1* KO cells restored normal gene expression (Figure 7E). Once again, *TREX1* KO cells did not express pro-inflammatory genes in the absence of chromosomal instability (Figure 7E). Taken together, these results indicate that ER-directed TREX1 localization and activity at ruptured MN is critical for inhibiting cGAS recognition of ruptured MN and the consequent pro-inflammatory transcriptional response.

## DISCUSSION

The findings reported herein identify TREX1-mediated partial degradation of micronuclear DNA as a novel mechanism of cGAS regulation in chromosomally unstable cells (Figure 7F). Our work also expands the list of TREX1 target substrates relevant to disease. The data are consistent with recent work showing that *TREX1* deletion enhances ISG expression in chromosomally unstable, BLM-deficient fibroblasts (Gratia et al., 2019). In contrast to the prevailing view of TREX1’s mechanism of action, we find that TREX1 is capable of restricting cGAS recognition of MN without completely eliminating MN or dsDNA from the cytosol. Instead, we find t hat T REX1 targets limited regions of micronuclear DNA for resection. It is tempting to speculate that TREX1 may preferentially target nucleosome-free DNA most likely to activate cGAS (Zierhut et al., 2019). Indeed prior studies have noted the resistance of chromatin to TREX1 digestion (Chowdhury et al., 2006). We further speculate that TREX1 degradation of micronuclear DNA is primed by a currently unknown endonuclease, but our data do not exclude other possibilities.

### Purification of ruptured MN

Despite established links to immune activation and chromosomal rearrangement, many aspects of ruptured MN have remained enigmatic and their structure and composition are largely unknown. These questions have persisted, in part, because of technical limitations: current methods separate MN from primary nuclei by centrifugation but cannot distinguish between ruptured and intact MN (Shimizu et al., 1996). Here, we report a novel method that uses genetically-encoded markers to distinguish and purify ruptured and intact MN. Using this technique, we have discovered that TREX1 stably associates with MN and resects micronuclear DNA after loss of micronuclear envelope integrity. We anticipate that our method will enable further insights into the consequences of micronuclear envelope rupture, including alterations to protein and DNA content.

### TREX1 damages chromosomal DNA to limit cGAS recognition of MN

Our data support a model in which TREX1 partially degrades micronuclear DNA to inhibit cGAS DNA sensing in chromosomally unstable cells (Figure 7F). Several pieces of evidence support a model of cGAS inhibition through direct TREX1 action at MN. First, cytosolic dsDNA was not increased after Mps1 inhibition or *TREX1* deletion. Second, MN-independent, cytosolic cGAS foci were not increased in chromosomally unstable or *TREX1* KO cells. Third, deleting the TREX1 C-terminus disrupted normal TREX1 localization to ruptured MN and compromised MN-associated DNA degradation and cGAS inhibition. Re-tethering TREX1ΔC to the ER via fusion to the Sec61-TMD restored cGAS inhibition, highlighting the importance of ER association for TREX1 function. These experiments further suggest retention of TREX1ΔC catalytic activity despite deletion of the C-terminus.

Damage to micronuclear DNA after micronuclear envelope rupture is associated with chromothripsis, but the causes of DNA damage at MN are poorly understood (Crasta et al., 2012; Hatch et al., 2013; Ly et al., 2017, 2019; Umbreit et al., 2020). Likewise, DNA bridges are another known precipitant of chromothripsis (Maciejowski et al., 2015; Maciejowski et al., 2019; Umbreit et al., 2020). Recent work indicates that TREX1 is dispensable for DNA bridge breakage and for copy number variations observed after spontaneous DNA bridge breakage in RPE1 cells (Umbreit et al., 2020). However, this study did not assess TREX1 localization to MN or DNA bridges and did not look for direct evidence of TREX1-dependent DNA damage at DNA bridges or ruptured MN. Our report is the first to identify an enzymatic source of DNA damage in MN after micronuclear envelope rupture. Further experimentation will be necessary to assess the extent of TREX1-dependent DNA damage in MN and DNA bridges and its role in chromothripsis.

Primary NE rupture is also associated with DNA damage (Denais et al., 2016; Irianto et al., 2017; Pfeifer et al., 2018; Raab et al., 2016; Xia et al., 2018). Our findings are consistent with recent work showing that TREX1 is the primary source of the DNA damage that occurs in the primary nucleus after compression-induced NE rupture (Nader et al., 2020). Consideration of this new data, together with the work presented here, suggests the existence of similar TREX1-dependent mechanisms underlying NE rupture-associated DNA damage at the primary nucleus and at MN.

### ER-tethering directs TREX1 substrate engagement

Frameshift mutations that remove the C-terminus of TREX1 do not compromise exonuclease activity but are nevertheless associated with autoimmune diseases such as retinal vasculopathy with cerebral leukodystrophy and systemic lupus erythematosus (Lee-Kirsch et al., 2007; Richards et al., 2007; de Silva et al., 2007). Previous studies have proposed that the TREX1 C-terminus possesses a distinct, nuclease-independent function in the regulation of ER oligosaccharyltransferase complex activity (Fermaintt et al., 2019; Hasan et al., 2015; Kucej et al., 2017; Yan, 2017). Consequently, deleting the TREX1 C-terminus results in autoimmune disease and inflammation via the accumulation of free glycans (Fermaintt et al., 2019; Hasan et al., 2015).

Our data show that the C-terminus of TREX1 also plays a critical role in promoting canonical TREX1 nuclease activity by promoting engagement with target DNA, such as MN, via ER-tethering. However, the TREX1 C-terminus appears to be dispensable for degrading transfected plasmid DNA, suggesting that TREX1 may be uniquely dependent on ER-tethering for degrading micronuclear DNA or other chromatin targets. The DNA substrates of TREX1 relevant to autoimmune disease have not been comprehensively characterized (Crow and Manel, 2015). Interestingly, our findings suggest that deletion of the TREX1 C-terminus may result in autoimmune disease due to ineffective engagement with critical targets, including MN, which can accumulate – albeit infrequently – in healthy tissue (Guo et al., 2019).

It is not currently clear how ER-tethering promotes TREX1 engagement with ruptured MN. The strong enrichment of TREX1 at ruptured MN indicates the presence of an active targeting mechanism. One possibility is that TREX1 is targeted to ruptured MN by mechanisms that normally promote NE repair. Repair of ruptured NEs is hypothesized to occur via the recruitment of new sheets of ER-derived membrane by cytosolic BAF (Denais et al., 2016; Halfmann et al., 2019; Young et al. 2020). BAF also recruits the ESCRT-III complex to reseal ruptured NEs (Denais et al., 2016; Halfmann et al., 2019; Raab et al., 2016). NE repair generally does not occur in MN (Hatch et al., 2013). However, BAF and ESCRT-III localize to MN after micronuclear envelope rupture (Liu et al., 2018; Vietri et al. 2019; Willan et al., 2019). Therefore, TREX1 localization to MN may depend on BAF-dependent recruitment of ER-associated membranes or remodeling of membranes by ESCRT-III.

## CONCLUSIONS

Chromosomally unstable tumor cells can adapt to chronic dsDNA in the cytosol by suppressing Type I interferon and activating non-canonical NF-κB (Bakhoum et al., 2018; Dou et al., 2017). Our work reveals that TREX1-mediated inhibition of cGAS sensing of ruptured MN may be one mechanism that chromosomally unstable tumor cells rely on to suppress anti-tumor immunity and cell autonomous growth inhibition resulting from chronic cGAS-STING pathway activation. Previous work has shown that TREX1 upregulation can inhibit radiotherapy-induced tumor immunogenicity (Vanpouille-Box et al., 2017). Our results suggest that TREX1 regulation of immunogenicity may be a more general phenomenon that extends to chromosomally unstable cancer cells. Therefore, TREX1-mediated DNA degradation in MN may enable cancer cells to benefit from chromosomal instability without the potentially lethal effects of cGAS-STING activation.

## ACKNOWLEDGEMENTS

We thank Titia de Lange, Sam Bakhoum, Philip Kranzusch, Wen Zhou, Christian Zierhut, Agnel Sfeir, and Zhijian Chen for advice and reagents, the Integrated Genomics Operation at MSKCC for assistance with Nanostring gene expression analysis, Matthieu Piel and Guilherme Nader for sharing unpublished data, and members of the Maciejowski lab for critical reading of this manuscript. Work in J.M.’s lab is supported by the NCI (R00CA212290), the Pew Charitable Trusts, the V Foundation, the Starr Cancer Consortium, the Emerald Foundation, the Geoffrey Beene and Ludwig Centers at MSKCC, and MSKCC core grant P30-CA008748.

## AUTHOR CONTRIBUTIONS

L.M. and J.M. designed the experiments. L.M. performed all of the cell biological experiments with help of K.C.. E.T. developed and performed experiments related to micronucleus purification. J.M. wrote the manuscript with the help of the other authors.

## METHODS

### Cell culture

MCF10A cells were from Maria Jasin (MSKCC) and cultured in 1:1 mixture of F12:DMEM media supplemented with 5% horse serum (Thermo Fisher Scientific), 2 0 n g/ml h uman E GF (Sigma), 0.5 mg/ml hydrocortisone (Sigma), 100 ng/ml cholera toxin (Sigma) and 10 μg/ml recombinant human insulin (Sigma). BT474 cells were cultured in RPMI 1640 supplemented with 10% FBS, 1% glucose, 1% sodium pyruvate. HCC1143 cells were cultured in RPMI 1640 supplemented with 10% FBS. RPE1-hTERT and MDA-MB-453 cells were cultured in F12:DMEM supplemented with 10% FBS. HEK293T, Phoenix, 293FT, HeLa, HT1080, CAL51, BT549, and U2OS were grown in DMEM supplemented with 10% FBS. All media was supplemented with 1% penicillin-streptomycin. Unless otherwise noted, all media and supplements were supplied by the MSKCC core facility.

For retroviral transduction, open reading frames were cloned into pQCXIN, pQCXIP, pQCXIB (Clontech), or pQCXIZ, which confer resistance to G418, puromycin, blasticidin, and zeocin. Constructs were transfected into Phoenix amphotropic packaging cells using calcium phosphate precipitation. Cell supernatants containing retrovirus were filtered, mixed 1:1 with target cell media and supplemented with 4 μg /ml polybrene. Successfully transduced cells were selected using G418 (Corning), puromycin (Fisher), blasticidin (Fisher), or zeocin (Life Technologies). Clones were isolated by limiting dilution or flow sorting.

For lentiviral transduction, open reading frames were cloned into pLenti CMV GFP Neo/Blast/Puro (Addgene 17447, 17445, 17448). Constructs were transfected into 293FT cells together with psPAX2 (Addgene 12260) and pMD2.G (Addgene 12259) using calcium phosphate precipitation. Supernatants containing lentivirus were filtered a nd supplemented with 4 μg/ml polybrene. Successfully transduced cells were selected as above.

### Immunofluorescence microscopy

For immunofluorescence microscopy of micronucleated cells, MCF10A cells were treated with 0.5 μM of the Mps1 inhibitor reversine for 72 h. Parental and 3×FLAG-TREX1 overexpressing HEK293T (+H2B-mCherry +GFP-cGAS) cells (Figure 1, S1, S4A,B,F,G) were treated for 48 h, while HeLa, RPE1 hTERT, HT1080, and U2OS cells (Figure 2F) were treated for 24h. For immunofluorescence staining for cGAS, Calreticulin, dsDNA, γH2AX, GFP, LSD1, RPA32 or TREX1, cells were carefully washed with PBS prior to fixation i n 2% paraformaldehyde i n PBS for 12 min. For staining of pIRF3, cells were incubated with TBS-TX (TBS supplemented with 0.1% Triton X-100) for 5 min and fixed with −20°C m ethanol and kept at −20°C for at least 1 h. Coverslips were washed with TBS, incubated in TBS with 0.5% Triton X-100 for 5 min and washed again with TBS. Coverslips were incubated in blocking buffer (1 mg/ml BSA, 3% goat serum, 0.1% Triton X-100, 1 mM EDTA in PBS) for 1 h and primary antibodies (see Key Resource Table), diluted in blocking buffer, were added for 2 h. After 4 washes with TBS-TX, coverslips were incubated with secondary antibodies (see Key Resource Table), diluted in blocking buffer, for 1 h, then washed 2 times with TBS-TX. DNA was stained with DAPI or Hoechst (both at 1 μg/ml) for 10 min, before coverslips were washed 2 times with TBS-TX and once with TBS. Coverslips were mounted in ProLong Gold Antifade Mountant (Life Technologies). Images were acquired on a DeltaVision Elite system equipped with a DV Elite CMOS camera, microtiter stage, and ultimate focus module (Z-stacks through the cells at 0.2 μm increments). For figures 6D, 7A, S7A images were processed by digital deconvolution of Z-stack image series using softWoRx software. Images were further edited with Adobe Photoshop CS5.1.

Quantification of RPA32-, γH2AX- or BrdU-positive MN was performed as follows: After deconvolution of Z-stack image series and maximum intensity projections, MN were identified by Hoechst signal. Then the LSD1 or cGAS signal was used to distinguish between intact (LSD-positive or cGAS-negative) and ruptured MN (LSD-negative or cGAS-positive). Finally MN were called RPA32 or γH2AX-positive, if there were more than 2 clear foci visible per MN.

For staining of cytosolic dsDNA (Figure S1E-H), cells were treated with TBS with 0.02% saponin for 5 min after fixation to selectively permeabilize the plasma membrane as previously described (Bakhoum et al., 2018).

### Native BrdU staining

For detection of ssDNA in MN, cells seeded on coverslips were preincubated with BrdU (10 μM) for 24 h, washed twice with media and treated with reversine (0.5 μM) for 48 h. Before BrdU and cGAS IF, cells were carefully washed with PBS, incubated in ice cold extraction buffer (10 mM Pipes-NaOH pH 7, 100 mM NaCl, 300 mM sucrose, 3 mM MgCl_2_, 1 mM EGTA, 0.5% Triton X-100) for 10 min. Fixation and staining steps, as well as quantification were performed as described above.

### Immunoblotting

For the Western blots shown in Figures S2A, S4C and S6A, cells were harvested by trypsinization and lysed in 1× Laemmli buffer (50 mM Tris, 10% glycerol, 2% SDS, 0.01% bromophenol blue, 2.5% β-mercaptoethanol) at 10^7^ cells/ml. Lysates were denatured at 95°C and DNA was sheared with a 28 ½ gauge insulin needle. Lysate equivalent to 1-2 × 10^5^ cells was resolved by SDS-PAGE (Life Technologies) and transferred to nitrocellulose membranes (Amersham). Membranes were blocked in 5% milk in TBS with 0.1% Tween-20 (TBS-T) and incubated with primary antibody (see Key Resource Table) overnight at 4°C, washed 4 times in TBS-T, and incubated for 1 h at room temperature with horseradish-peroxidase-conjugated secondary antibody (see Key Resource Table). After four washes in TBS-T, membranes were rinsed in TBS and imaging was performed using enhanced chemiluminescence (Thermo Fisher).

For quantitative Western blotting (Figures 6A, S1D, S3A and S6G), cells were lysed in RIPA buffer 25 mM Tris-HCl pH 7.6, 150 mM NaCl, 1% NP-40, 1% sodium deoxycholate, 0.1% SDS), supplemented with phosphatase inhibitors (10 mM NaF, 20 mM β-glycerophosphate) and protease inhibitor (Thermo Scientific) at 10^7^ cells/ml and incubated on ice for 30 min. After centrifugation (21,000 × g, 30 min, 4°C), protein concentration of the supernatant was determined using BCA protein assay (Thermo Fisher) and 50 μg (1C and S2F) or 20 μg (S3D, S6B and S7A) protein was loaded per sample. Membranes were blocked in Odyssey blocking buffer in TBS (Li-COR). Primary antibodies were diluted in blocking buffer supplemented with 0.2% Tween and incubated with membranes overnight at 4°C. Secondary antibodies (Alexa Fluor 680 and 800; Key Resource Table) were used at 1:20,000 dilutions in blocking buffer supplemented with 0.2% Tween. Fluorescence was measured using an infrared imaging scanner (Odyssey; LI-COR) according to the manufacturer’s instructions.

For Western blots of purified M N (Figures 1G; S1A,B), samples were lysed in 1× Laemmli buffer at 2.5 × 10^7^ MN/ml. Lysates were denatured at 95°C and DNA was sonicated using a Bioruptor 300 (Diagenode) on the high setting for 8 cycles 30 sec ON / 30 sec OFF. Lysate equivalents to 0.5×10^6^ MN were loaded in each lane and processed as described above for quantitative Western blotting.

### Gene expression analysis

Total RNA was isolated from 2 × 10^6^ cells using the Quick-RNA miniprep kit (Zymo) according to the manufacturer’s instructions.

cDNA was generated from 500 ng RNA using the SuperScript IV first-strand s ynthesis s ystem (Life Technologies) with random hexamer priming. qPCR was performed with gene specific primers (see Key Resource Table) using Sybr green detection on a Light Cycler 480 (Roche) or Quantstudio6 (Applied Biosystems) cycler. Relative transcription levels were calculated by normalizing to ACT expression.

NanoString was used to directly quantify mRNA transcripts. Isolated RNA was hybridized with reporter and capture probes of the nCounter Human Inflammation V2 Panel (Figure 6F and S6I) or a custom made nCounter gene expression code set (Figure 7E, see Key Resource Table) according to the manufacturer’s instructions (NanoString Technologies). Data was analyzed using nSolver Analysis software.

### 2′3′-cGAMP quantification

4 × 10^6^ of MCF10A or M2p1 cells were seeded into 15-cm dishes, and 24 h later cells were treated with reversine (0.5 μM) or doxycycline (1 μg/ml), respectively. For stimulation with dsDNA, 1.5 × 10^6^ of MCF10A cells were seeded into 10-cm dishes, and 24 h later transfected with 4 μg of pMaxGFP plasmid (Lonza) using Fugene HD transfection reagent (Promega) per manufacturer’s instructions. 72 h after reversine and doxycycline addition or 24 h after transfection, cells were harvested, washed with PBS, pelleted, and stored at −80°C. To quantify 2′3′-cGAMP levels, 8 × 10^6^ cells (reversine) or 2 × 10^6^ cells (plasmid DNA) were thoroughly resuspended in 120 μL lysis buffer (20 mM Tris-HCl pH 7.7, 100 mM NaCl, 10 mM NaF, 20 mM β-glycerophosphate, 5 mM MgCl2, 0.1% Triton X-100, 5% glycerol) and lysed with a 28 ½ gauge needle. Lysates were incubated on ice for 30 min, centrifuged at 16,000 × g, 4°C for 10 min and 2′3′-cGAMP levels were quantified using the 2′3′-cGAMP ELISA Kit (Arbor Assays) according to the manufacturer’s instructions.

### Live-cell imaging

Cells were plated onto 35 mm glass bottom dishes (Cellvis) 48 h before imaging. One h before imaging, media was replaced with fresh medium. Live-cell imaging was performed at 37°C and 5% CO2 using a Nikon Eclipse Ti2-E equipped with a CSU-W1 spinning disk with Borealis microadapter, Perfect Focus 4, motorized turret and encoded stage, polycarbonate thermal box, 5 line laser launch [405 (100 mw), 445 (45 mw), 488 (100 mw), 561 (80 mw), 640 (75 mw)], PRIME 95B Monochrome Digital Camera, and environmental enclosure (Tokai Hit). Objective lenses included CFI Plan Apo Lambda 40x 0.95 NA and CI Plan Apo Lambda 60x 1.40 NA. Images were acquired using NIS-Elements Advanced Research Software on a Dual Xeon Imaging workstation. Maximum intensity projection of z-stacks and adjustment of brightness and contrast were performed using Fiji software. Images were cropped and assembled into figures using Photoshop CS5.1 (Adobe).

### MN purification

MN purification protocol was adapted from a previously described method (Shimizu et al., 1996). Parental and 3xFLAG-TREX1 overexpressing HEK293T (+H2B-mCherry +GFP-cGAS) cells were treated with 0.5 μM reversine 48 h prior to harvesting 10^8^-10^9^ cells were harvested and washed twice in DMEM without serum. Washed cells were resuspended in pre-warmed (37°C) DMEM without serum supplemented with cytochalasin B (Cayman) at 10 μg/ml at a concentration of 10^7^ cells/ml DMEM and incubated at 37°C for 30 minutes. Cells were centrifuged at 300 x g for 5 minutes and cell pellet was resuspended in cold lysis buffer (10 mM Tris-HCl, 2 mM Mg-acetate, 3 mM CaCl2, 0.32 M sucrose, 0.1 mM EDTA, 0.1% (v/v) NP-40, pH 8.5) freshly complemented (with 1 mM dithiothreitol, 0.15 mM spermine, 0.75 mM spermidine, 10 μg/ml cytochalasin B and protease inhibitors) at a concentration of 2 × 10^7^ cells/ml lysis buffer. Resuspended cells were then dounce homogenized by 10 strokes with a loose-fitting pestle. At this point a verification step was performed by mixing a small volume of lysed cells with an equal volume of PBS + 0.5 μg/ml DAPI to confirm M N r elease by fluorescence m icroscopy. Cell lysates were then mixed with an equal volume of ice cold 1.8 M sucrose buffer (10 mM Tris-HCl, 1.8 M sucrose, 5 mM Mg-acetate, 0.1 mM EDTA, pH 8.0) freshly complemented (with 1 mM dithiothreitol, 0.3% BSA, 0.15 mM spermine, 0.75 mM spermidine) before use. 10 ml of this mixture (lysed cells + 1.8 M sucrose buffer) was then layered on top of a two-layer sucrose gradient (prepared by slowly adding 20 ml of 1.8 M sucrose buffer slowly on top of 15 ml 1.6 M sucrose buffer in a 50 ml conical tube). This mixture was then centrifuged in a JS-5.2 swinging bucket rotor (Beckman) at 944 × g for 20 min at 4°C. Generally, fractions were collected as follows: upper 2 ml contain debris and is discarded; next 5–6 ml contains MN and is collected; final 38 m l contains primary nuclei and is discarded. At this point a verification s tep w as often included to most accurately identify fractions containing MN and lacking debris and primary nuclei. Fractions containing MN were then pooled and diluted 1:5 with FACS buffer (ice cold PBS supplemented with 0.3% BSA, 0.1% NP-40 and protease inhibitors). Diluted MN were then centrifuged at 944 × g in JS-5.2 swinging bucket rotor for 20 min at 4°C. Supernatant was then removed by aspiration and MN were resuspended in 2–4 ml of FACS buffer supplemented with 2 μg/ml DAPI. Resuspended samples were filtered t hrough a 4 0 μm ministrainer (PluriSelect) into FACS tubes. MN were then sorted by FACSAria (BD Biosciences) into FACS buffer at the MSKCC Flow Cytometry Core Facility. Default FSC and DAPI thresholds were lowered and a log scale was used to visualize MN. Sorted MN were centrifuged at 4000 × g in JS-5.2 swinging bucket rotor for 20 min at 4º C and the pellets were stored at −80°C before lysis for Western blotting or used directly for immunofluorescence microscopy.

### Immunofluorescence of purified MN

For immunofluorescence m icroscopy o f purified MN, samples were fixed in 2 % paraformaldehyde for 10 minutes by adding the appropriate volume of 16% paraformaldehyde to the samples collected in FACS buffer (ice cold PBS supplemented with 0.3% BSA, 0.1% NP-40 and protease inhibitors). Then, samples were loaded in cytofunnels (Thermo Fisher), centrifuged on microscopy slides using a cytospin 4 centrifuge (Thermo Fisher) and dried overnight at room temperature. Each sample was circled with an aqua-hold pap pen (Fisher Scientific) prior to incubation in blocking buffer, antibody and DAPI staining (see above).

## SUPPLEMENTAL FIGURES

**Figure S1.**
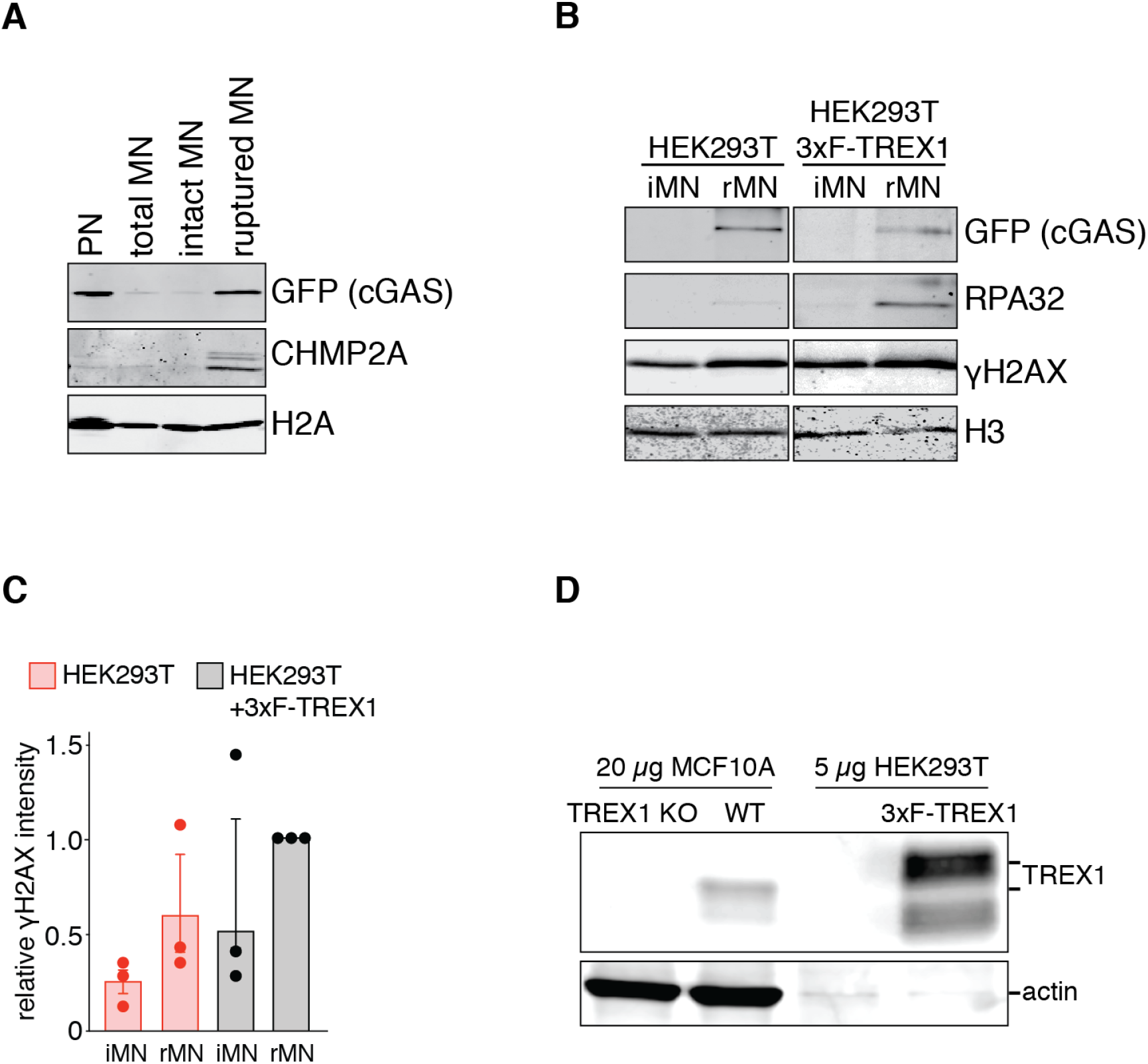
TREX1 overexpression in HEK293T cells. Related to Figure 1. (A) Immunoblotting for the indicated proteins in primary nuclei (PN), ruptured and intact MN sorted from HEK293T (+H2B-mCherry +GFP-cGAS) cells. (B) Immunoblotting for the indicated proteins in ruptured and intact MN sorted from parental and 3xFLAG-TREX1 overexpressing HEK293T (+H2B-mCherry +GFP-cGAS) cells. (C) Quantification of relative γH2AX signals normalized to loading control (SMC1 or histone H2A) as shown in (B). Mean and s.d. of *n* = 3 independent biological replicates are shown. (D) Immunoblotting for TREX1 and actin in MCF10A (wt and *TREX1* KO) as well as parental and 3xFLAG-TREX1 overexpressing HEK293T (+H2B-mCherry +GFP-cGAS) cells.

**Figure S2.**
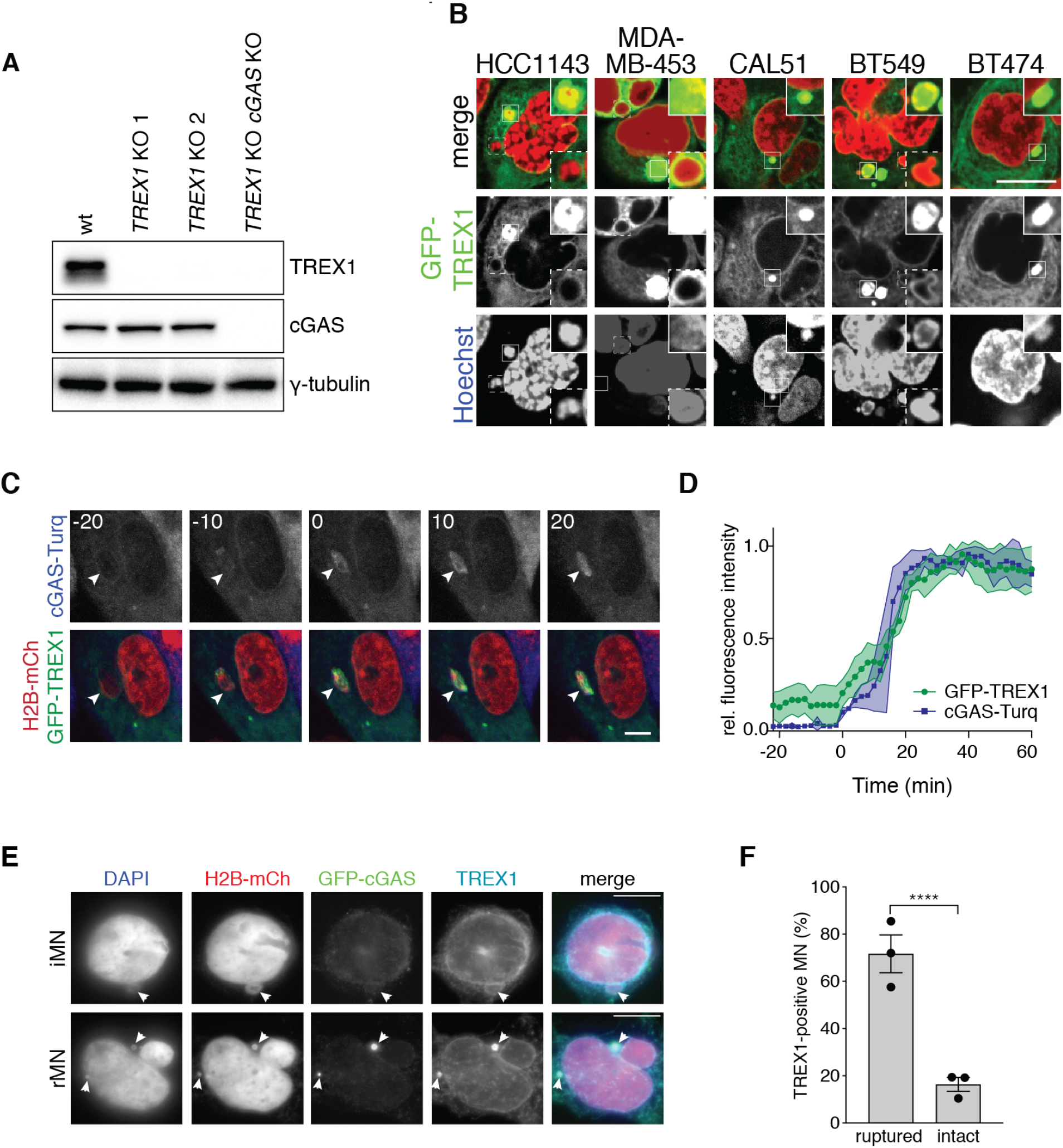
TREX1 localizes to MN and DNA bridges. Related to Figure 2. (A) Immunoblotting for TREX1, cGAS, and γ-tubulin in the indicated MCF10A cells. (B) TREX1 localizes to a fraction of MN in chromosomally unstable human breast cancer cell lines. Immunofluorescence for TREX1 in the indicated cell lines. DNA was stained with Hoechst 33342. Insets show magnified regions (marked by white boxes) to highlight TREX1 localization to MN. (C) Frames from live-cell imaging of Mps1i-treated MCF10A cells expressing GFP-TREX1, H2B-mCherry and mTurquoise2-cGAS to assay for micronuclear envelope rupturing (t = 0, micronuclear envelope rupture). (D) Quantification of the relative fluorescence signal intensities of GFP-TREX1 and mTurquoise2-cGAS as shown in (C). Mean and s.d. of *n* = 8 MN are shown. (E) Immunofluorescence for GFP and TREX1in 3xFLAG-TREX1 overexpressing HEK293T (+H2B-mCherry +GFP-cGAS) cells treated with Mps1i for 48 h. DNA was stained with DAPI. GFP (cGAS) staining serves as a marker for ruptured MN. Arrowheads mark MN. (F) Quantification of the percentage of TREX1-positive ruptured and intact MN as shown in (E). Mean and s.d. from *n* = 3 independent biological replicates are shown (>130 MN quantified per replicate and cell line). *P* values were calculated by Student’s t-test (\*\*\*\**P* < 0.0001). All scale bars = 10 μm.

**Figure S3.**
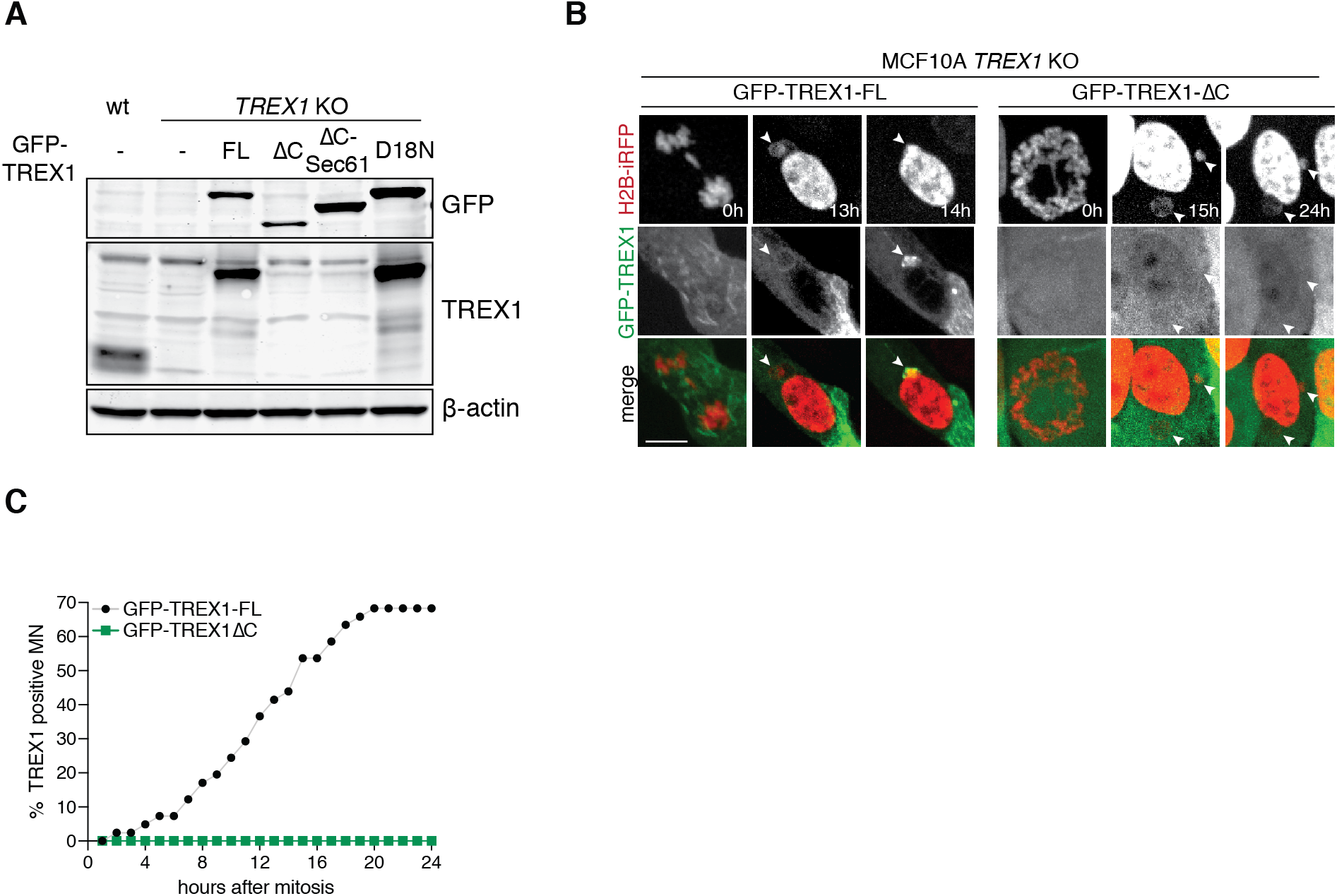
Loss of TREX1 C-terminus disrupts localization to MN. Related to Figure 3. (A) Immunoblotting for GFP, TREX1 and β-actin in the indicated MCF10A cell lines, overexpressing GFP-TREX1 transgenes, -FL (full length), ΔC (TREX1 truncation, aa 1-235), ΔC-Sec61 (chimeric protein consisting of TREX1 aa1-235 fused to the Sec61-TMD) and D18N (nuclease deficient TREX1). Note that the antibody used here to detect TREX1 is raised against the C-terminus and therefore doesn’t recognize the C-terminally truncated variants of TREX1 (Abcam; see methods). (B) Frames from live-cell imaging of Mps1i-treated MCF10A *TREX1* KO cells expressing GFP-TREX-FL or ΔC and H2B-iRFP to mark chromatin. Time after mitotic exit is marked in the bottom right corner. Arrowheads mark MN. Scale bar = 10 μm (C) Quantification of the cumulative percentage of TREX1 positive MN as shown in (B). MN were followed from mitotic exit until the subsequent mitosis or for a max of 24 h. *n* = 2 independent biological replicates with 45 MN per cell line and replicate.

**Figure S4.**
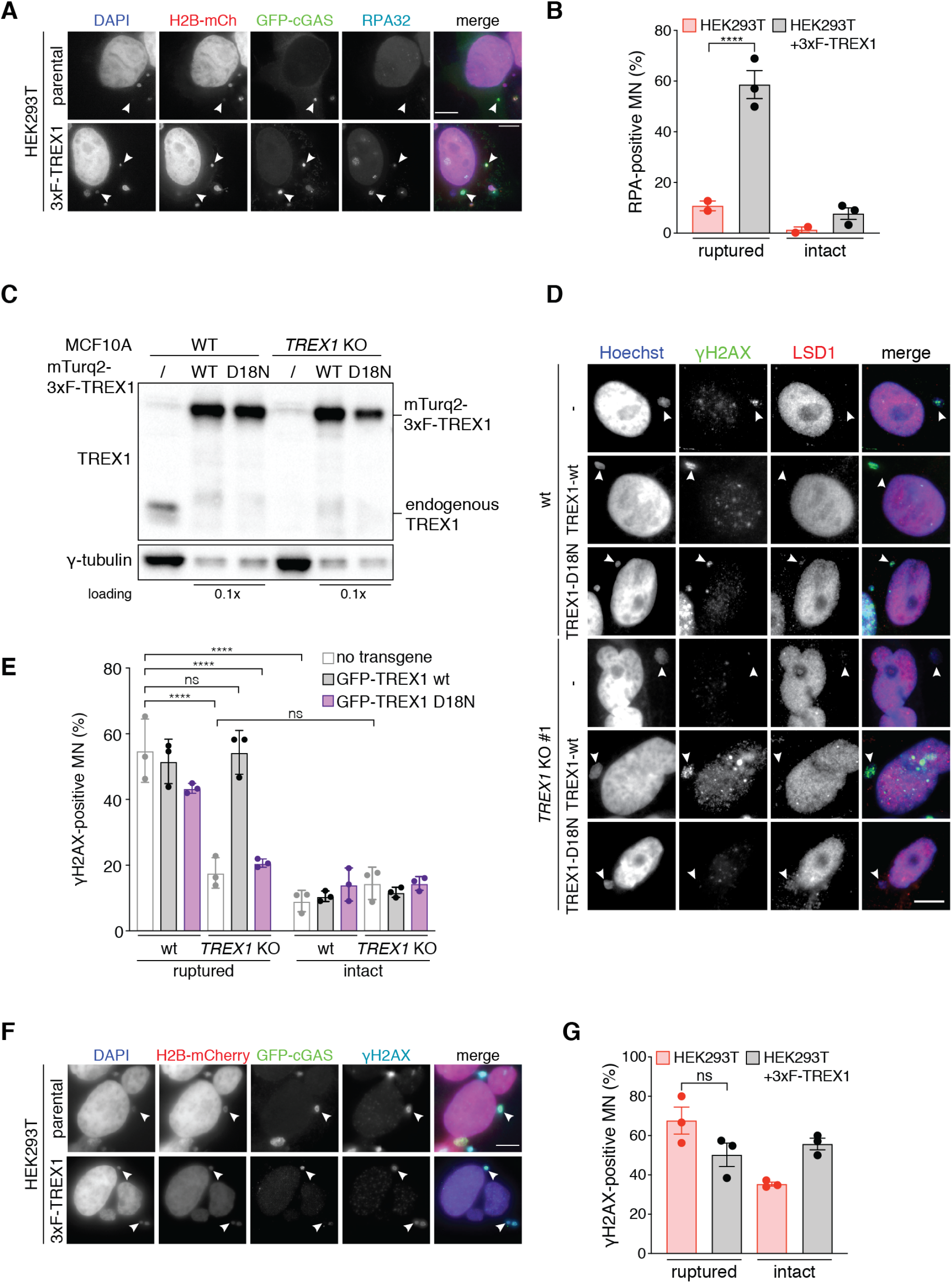
TREX1 causes DNA damage in ruptured MN. Related to Figure 4. (A) Indicated HEK293T (+H2B-mCherry +GFP-cGAS) cells were treated with Mps1i for 48 h. Immunofluorescence for RPA32 and GFP. DNA was stained with DAPI. cGAS signal is used to infer micronuclear envelope rupture. Arrowheads mark ruptured MN. (B) Quantification of the percentage of RPA32-positive ruptured and intact MN as shown in (A). Mean and s.d. from *n* = 3 independent biological replicates are shown (>200 MN quantified per replicate and cell line). *P* values were calculated by Student’s t-test (\*\*\*\*\**P* < 0.0001). (C) Immunoblotting for TREX1 and γ-tubulin in the indicated MCF10A cell lines (wt and *TREX1* KO 1), overexpressing mTurq2-3xFLAG-TREX1-wt or D18N (nuclease deficient). (D) Immunofluorescence for γH2AX and LSD1 in the indicated MCF10A cells. DNA was stained with Hoechst 33342. Scale bar = 10 μm. Arrowheads mark ruptured MN. (E) Quantification of the percentage of γH2AX-positive ruptured and intact MN as shown in (D). Mean and s.d. from *n* = 3 independent biological replicates are shown (>200 MN quantified per replicate and cell line). *P* values were calculated by two-way ANOVA with Sidak’s multiple comparisons test (\*\*\*\**P* < 0.001). (F) Indicated HEK293T (+H2B-mCherry +GFP-cGAS) cells were treated with Mps1i for 48 h. Immunofluorescence for γH2AX and GFP. DNA was stained with DAPI. cGAS signal is used to infer micronuclear envelope rupture. Arrowheads mark ruptured MN. (G) Quantification of the percentage of γH2AX-positive ruptured and intact MN as shown in (F). Mean and s.d. from *n* = 3 independent biological replicates are shown (>130 MN quantified per replicate and cell line). *P* values were calculated by Student’s t-test (ns = not significant). All scale bars = 10 μm.

**Figure S5.**
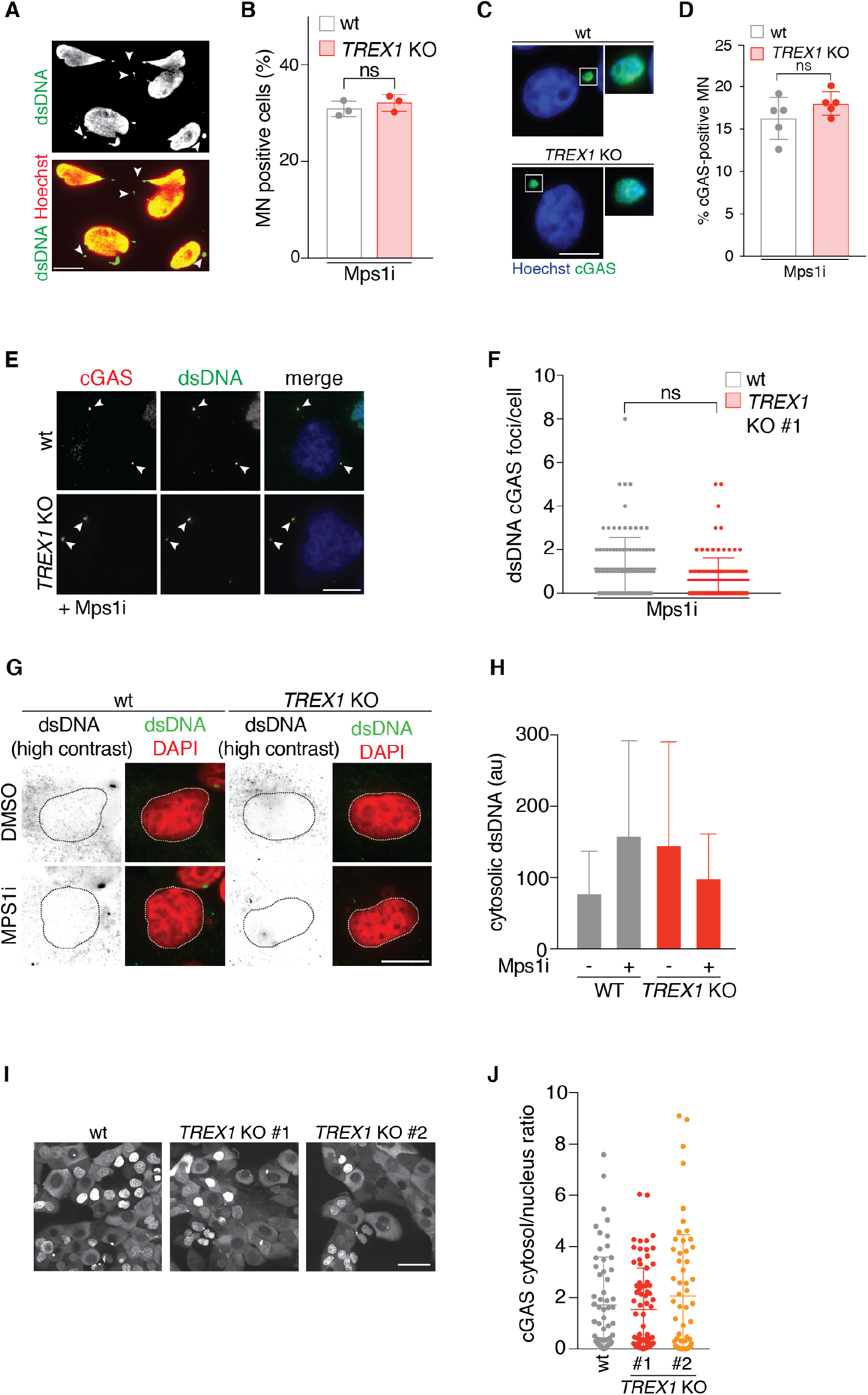
Mps1i treatment induces equivalent levels of chromosomal instability in MCF10A wt and *TREX1* KO cells. Related to Figure 5. (A) Immunofluorescence for dsDNA in MCF10A cells 72 hours after treatment with Mps1 inhibitor (Mps1i = reversine). Arrowheads indicate MN and DNA bridges. Scale bar = 10 μm. (B) Quantification of the percentage of cells with MN in (A). Mean and s.d. of *n* = 3 independent biological replicates are shown (>200 cells quantified per replicate and cell line). *P* values were calculated by Student’s t-test (ns = not significant). (C) TREX1 deficiency does not disrupt cGAS localization to MN. Immunofluorescence for cGAS of the indicated MCF10A cells 72 h after treatment with Mps1i. (D) Quantification of the percentage of cGAS positive MN in (C). Mean and s.d. of *n* = 5 independent biological replicates are shown (>220 MN quantified per replicate and cell line). *P* values were calculated by Student’s t-test (ns = not significant). (E) Immunofluorescence for cGAS and dsDNA in the indicated MCF10A cells 72 h after treatment with Mps1i. Cells are fixed and permeabilized using a method for selective plasma membrane permeabilization (see methods). DNA was stained with Hoechst 33342. Arrowheads indicate cytosolic foci marked by cGAS and dsDNA. Scale bar = 10 μm. (F) Quantification of the number of cGAS and dsDNA double positive foci per cell as shown in (E). Mean and s.d. of *n* = 1 biological replicates are shown (>185 cells quantified per replicate and cell line). *P* value was calculated by Student’s t-test (ns = not significant). (G) Immunofluorescence for dsDNA in the indicated MCF10A cells 72 h after treatment with DMSO or Mps1i. Cells were fixed and permeabilized using a method for selective plasma membrane permeabilization (see methods). DNA was stained with Hoechst 33342. Scale bar = 10 μm. dsDNA images are inverted and highly contrasted to show minimal dsDNA signal in the cytosol, dashed circle marks location of primary nucleus. (H) Quantification of cytosolic dsDNA signal intensity as shown in (G). Mean and s.d. from *n* = 2 independent biological replicates are shown. At least 200 cells per condition were imaged and quantified. (I) TREX1 deficiency does not impact the distribution of GFP-cGAS between the nucleus and the cytosol. Live-cell images of GFP-cGAS in the indicated MCF10A cells. Scale bar = 20 μm. (J) Quantification of the ratio of cGAS signal intensity between the cytosol and nucleus as shown in (I). Mean and s.d. of individual cell data points from *n* = 1 experiment (>60 cells per replicate and cell line) are shown.

**Figure S6.**
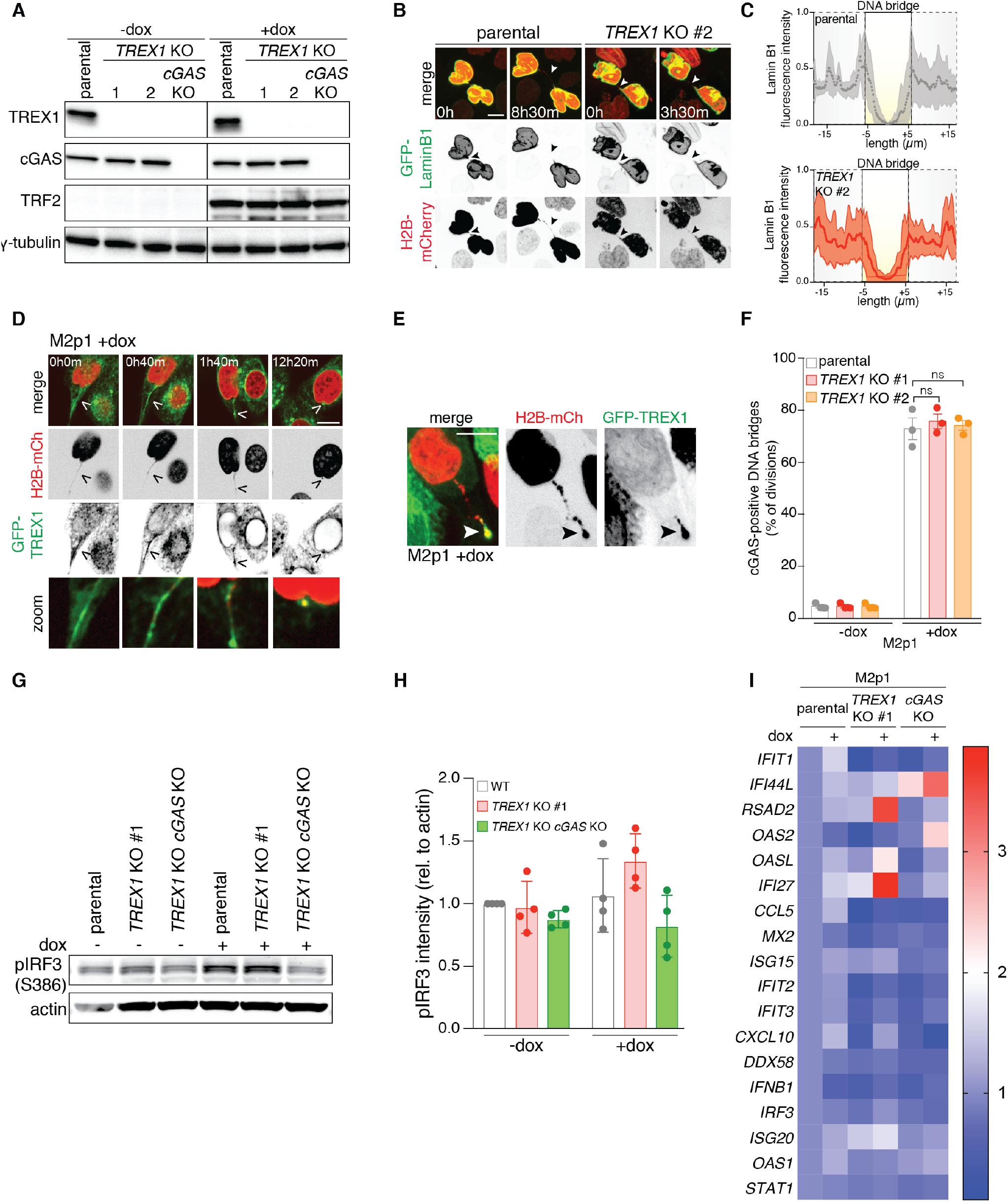
TREX1 inhibits cGAS sensing of DNA bridges. Related to Figure 5 and 6. (A) Immunoblotting for TREX1, cGAS, dominant negative TRF2 (TRF2-DN) and γ-tubulin in the indicated M2p1 (MCF10A *TP53* KO doxi::TRF2-DN) cells 72 h after DMSO or doxycycline treatment. (B) TREX1 deficiency does not influence loss of LaminB1 at DNA bridges. Frames from live-cell imaging of doxycycline-treated M2p1 (MCF10A *TP53* KO doxi::TRF2-DN) cells expressing GFP-LaminB1 and H2B-mCherry to mark chromatin. Relative time between images is marked in hours and minutes in the bottom left of each panel. Scale bar = 10 μm. Arrowheads mark DNA bridges. (C) Quantification of LaminB1 at DNA bridges. Mean and s.d. of *n* = 10 DNA bridges as shown in (B). (D,E) Frames from live-cell imaging of M2p1 cells expressing GFP-TREX1 and H2B-mCherry to mark chromatin treated with doxycycline to induce DNA bridge formation. Relative time between images is marked in hours and minutes in the top left of each panel. Arrowheads mark nuclei connected by a DNA bridge. (E) Example of GFP-TREX1 localization to a broken DNA bridge fragment. Arrowheads mark DNA bridges. (F) TREX1 deficiency does not disrupt cGAS localization to DNA bridges. Quantification of the number of cGAS positive bridges as a percentage of cell divisions as shown in (Figure 5H). Mean and s.d. of *n* = 3 independent biological replicates are shown. >50 cells were analyzed per condition per experiment. *P* values were calculated by one-way ANOVA with Tukey’s multiple comparisons test (ns = not significant). (G) Immunoblotting for IRF3 S386 phosphorylation and actin in the indicated M2p1 cells 72 h after DMSO or doxycycline treatment. (H) Quantification of pIRF3(S386) relative to corresponding actin signal as shown in (G). Mean and s.d. of *n* = 3 independent biological replicates are shown. (I) Heat map of Nanostring analysis of gene expression in the indicated M2p1 cells. RNA was isolated 72 h after DMSO or doxycycline treatment and Nanostring analysis was performed using a custom CodeSet (see methods). All scale bars = 10 μm.

**Figure S7.**
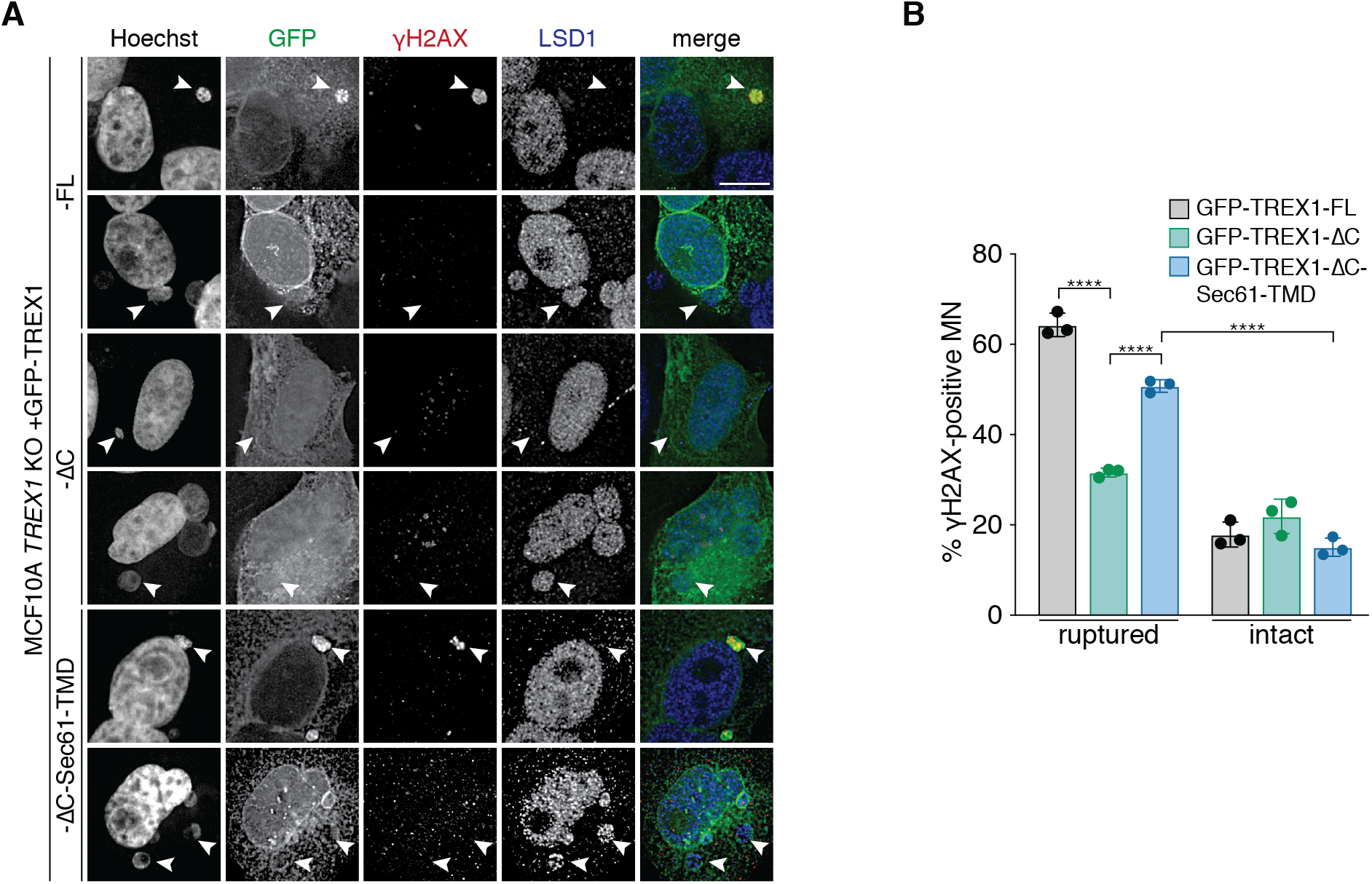
DSBs in ruptured MN are dependent on ER-tethering of TREX1. Related to Figure 7. (A) Immunofluorescence for γH2AX and LSD1 in the MCF10A *TREX1* KO cells expressing GFP-TREX1-FL (full length), ΔC (aa1-235) and ΔC-Sec61-TMD. DNA was stained with Hoechst 33342. Merge shows GFP, γH2AX and LSD1. Scale bar = 10 μm. Arrowheads mark ruptured (white) and intact (red) MN. (B) Quantification of the percentage of γH2AX-positive ruptured and intact MN as shown in (A). Mean and s.d. from *n* = 3 independent biological replicates are shown (>150 MN quantified per replicate and cell line). *P* values were calculated by two-way ANOVA with Sidak’s multiple comparisons test (\*\*\*\**P* < 0.0001).

